# Hindlimb kinematics, kinetics, and muscle dynamics during sit-to-stand and sit-to-walk transitions in emus (*Dromaius novaehollandiae*)

**DOI:** 10.1101/2024.02.14.580291

**Authors:** Yuting Lin, Jeffery W. Rankin, Luís P. Lamas, Mehran Moazen, John R. Hutchinson

**Affiliations:** Structure & Motion Laboratory, Department of Comparative Biomedical Sciences, Royal Veterinary College, Hatfield, United Kingdom; Pathokinesiology Laboratory, Rancho Los Amigos National Rehabilitation Center, Downey, CA, United States; CIISA, Faculty of Veterinary Medicine, University of Lisbon, Portugal; Department of Mechanical Engineering, University College London, London, UK

**Keywords:** emu, sit-to-stand, sit-to-walk, musculoskeletal model, inverse dynamics, static optimisation

## Abstract

Animals not only need to walk and run but also lie prone to rest and then stand up. Sit-to-stand (STS) and sit-to-walk (STW) transitions are vital behaviours little studied in species other than humans so far, but likely impose biomechanical constraints on limb design because they involve near-maximal excursions of limb joints that should require large length changes and force production from muscles. By integrating data from experiments into musculoskeletal simulations, we analysed joint motions, ground reaction forces, and muscle dynamics during the STS and STW transitions in a large terrestrial, bipedal, and cursorial bird: the emu (*Dromaius novaehollandiae*, ∼ 30 kg). Simulation results suggest that in both STS and STW, emus operate near the functional limits (∼ 50 % of shortening/lengthening) of some of their hindlimb muscles, particularly in distal muscles with limited capacity for length change and leverage. Both movements involved high muscle activations (> 50 %) and force generation of major joint extensor muscles early in the transition. STW also required larger net joint moments and non-sagittal motions, entailing greater demands for muscle capacity. Whilst our study involves multiple assumptions, our findings lay the groundwork for future studies to understand, for example, how tendon contributions may reduce excessive muscle demands, especially in the distal hindlimb. As the first investigation into how an avian species stands up, this study provides a foundational framework for future comparative studies investigating organismal morphofunctional specialisations and evolution, offering potential robotics and animal welfare applications.

**Summary statement:** The study demonstrates the dynamics, biomechanical constraints, and musculotendinous control strategies during the sit-to-stand/walk transitions for a large bipedal bird – the emu – informing morphology, evolution, and potential robotic applications.

## Introduction

The ability to perform sit-to-stand (STS) and sit-to-walk (STW) behaviours are fundamental for humans (e.g., Aissaoui and Dansereau, 1999; Sloot *et al*., 2020; Smith, Reilly and Bull, 2020) and animals (e.g., Lidfors, 1989; Gardner, 2011; Ellis et al., 2018; Brouwers *et al*., 2023). These behaviours must overcome gravitational constraints to substantially elevate the body’s centre of mass (COM) from a flexed initial limb posture, likely resulting in large joint moments and potentially unfavourable effective mechanical advantage (EMA) (Biewener, 1989). In humans, both STS and STW require considerable muscle strength and coordination for task execution and balance control (Ellis *et al*., 1984; Schultz *et al*., 1992; Doorenbosch *et al*., 1994; Roebroeck *et al*., 1994; Riley *et al*., 1997; Dehail *et al*., 2007; Yoshioka *et al*., 2009), or older adults, these activities even approach the upper limits of muscle capacity (Hughes, 1996; Hortobágyi *et al*., 2003). However, despite extensive studies on movement patterns and muscle engagement (Hughes *et al*., 1994; Smith *et al*., 2020; Perera *et al*., 2023), the control strategies used during these movements remain elusive even for humans (e.g., Pandy, Garner and Anderson, 1995; Bobbert *et al*., 2016; Actis *et al*., 2018; Shia *et al*., 2018). Remarkably, research on biomechanics of STS and STW in animals is extremely scarce, with only three studies on dogs as examples (Feeney *et al*., 2007; Ellis *et al*., 2018; Triviño *et al*., 2023),.

Birds, especially large, cursorial species including ostriches and emus, present a unique opportunity to understand limb structure and locomotor function (e.g., Carrano, 1999). Large, cursorial bird species are known for their remarkable speeds and efficient locomotion. Ostriches, for example can achieve extraordinary speeds (greater than 50 km h^-1^) and have a remarkable economy of locomotion due to elevated storage and release of elastic energy in tendons, with muscle fibres predicted to act either approximately isometric or slowly shortening (Rubenson *et al*., 2011; Smith and Wilson, 2013; Rankin, Rubenson and Hutchinson, 2016). These simulated fibre actions during locomotion are also consistent with studies of *in vivo* muscle function in other species (e.g., Biewener, 1998; Roberts *et al*., 1998; Fukunaga *et al*., 2001; Daley and Biewener, 2003; Lichtwark and Wilson, 2006; Lichtwark, Bougoulias and Wilson, 2007). However, when compared to other forms of locomotion, STS and STW impose unique musculoskeletal demands due to potentially large joint moments with a low strength-to-weight ratio (Biewener, 1989). In particular, the challenges faced by cursorial species during STS and STW, including substantial fibre length change and force production, likely are compounded by their elongated, flexed limbs and specialised muscular configurations (e.g., allometrically shorter muscle fibres in distal limbs) (Maloiy *et al*., 1979; Biewener, 2005; Lamas et al., 2014; Dick and Clemente, 2017; Bishop, Wright and Pierce, 2021). Understanding how these cursorial birds stand up should provide valuable insights into how bipeds other than humans overcome the challenges of STS and STW.

This study investigates the movement dynamics, biomechanical constraints, and musculotendinous control strategies during the STS and STW transitions in emus (*Dromaius novaehollandiae*). Emus serve as an ideal avian model due to their cursorial adaptations, manageability, and limb structure, offering a compelling comparative basis to humans. Our objectives are twofold: firstly, to quantify the patterns of hindlimb kinematics and kinetics in emu STS and STW behaviours, and secondly, to identify the mechanical constraints and possible strategies used by emus in performing the two tasks. We hypothesised that: (i) both transitions would require high muscle activations in the key hindlimb extensor muscles and muscles whose primary actions are non-parasagittal; (ii) emu hindlimb muscles would operate near their functional limits, especially for distal muscles; (iii) hindlimb tendons would play important roles in preventing large muscle fibre forces, activations, and length changes during transitions; (iv) STW in emus would entail greater demands for muscle capacity than STS, because it should involve greater forces and ranges of motion.

## Materials and methods

### Overview

The study comprises four primary stages, described in detail below. First, we collected STS and STW kinematic and kinetic data from two subadult male emus (*Dromaius novaehollandiae*; body mass ∼ 30 kg). We then combined these empirical data into a detailed musculoskeletal model of an emu hindlimb to estimate hindlimb net joint moments using OpenSim’s inverse dynamics routine (Delp *et al*., 2007). For comparative purposes, we focused primarily on the ‘flexion momentum’ and ‘ascending’ phases (Kralj, Jaeger and Munih, 1990; Schenkman *et al*., 1990; and see below) of both movements – from the onset of the movements until standing upright or initiating gait. Following this, we used the calculated joint moments to determine the required muscle activations, normalised fibre lengths, and muscle forces using OpenSim’s static optimisation routine (Delp *et al*., 2007), accompanied by sensitivity analyses of tendon slack lengths (TSLs). In the static optimisation simulations, we assumed rigid tendons to test whether muscle fibres alone could meet the joint moment requirements. Finally, to assess the roles of tendons, we conducted dynamic simulations using OpenSim Moco incorporating non-rigid tendon characteristics and passive muscle fibre force generation (Dembia *et al*., 2020). Detailed procedures regarding the collection of experimental data, development of the musculoskeletal model, and application of optimisation frameworks are expanded upon below.

### Animals

One emu cadaver (male; body mass 48.8 kg) was used for musculoskeletal model development (Lamas, 2015). Three emus were trained to complete the STS and STW tasks; however, only two of the birds were compliant with the procedures necessary for data collection and analysis. All emus were hatched at a commercial breeding farm in the United Kingdom and subsequently reared from four weeks of age at the Royal Veterinary College. Their diet consisted primarily of a commercial ostrich pelleted diet supplemented with grass. From six weeks of age, they had unrestricted access to both commercial food and grass. At 24 weeks, their diet transitioned from an ostrich grower diet to adult ostrich pelleted food (Dodson and Horrel Ltd., Kettering, Northamptonshire, UK). Throughout their development, no constraints were placed on their regular exercise regimen, and all emus had equal access to exercise areas and conditions. Approval for all studies involving these animals was obtained from the Royal Veterinary College’s Ethics and Welfare Committee, following the Animals (Scientific Procedures) Act 1986 under a Home Office license number 70/7122 to Alan Wilson.

### Experimental data

Data collection occurred indoors at the Royal Veterinary College’s Structure and Motion Lab. The two subadult male emus used in data collection weighed ∼ 27.4 kg and ∼ 28.5 kg. In all trials, the birds started in a near-symmetrical, crouched position. We classified trials where minimal forward movement occurred upon standing upright as a STS transition according to Riley *et al*., (1991), and defined trials that involved at least one complete stride forward during rising as a STW transition per Perera *et al*., (2023). We defined the STS transition using three primary phases marked by events based on human STS phase definitions (Kralj *et al*., 1990; Schenkman *et al*., 1990) (**Figure 1C**): (i) flexion momentum phase: this phase began at movement onset (identified by a 5 % change in the vertical ground reaction force (GRF)) and concluded when the tarsometatarsus lifted off from the ground (marked by a 5 mm change in the ankle marker height); (ii) ascending phase: started at heel-off and continued until minimum variation was observed in the vertical and cranial-caudal GRF (determined via the root mean squared method); (iii) stabilisation phase: started at the end of the ascending phase and ended when the vertical GRF equalled body weight and fluctuations of 0.1 % – characteristic of quiet standing – were detected. STW in emus was a fluid transition from STS to gait. To compare between the two movements, we defined the first two phases of the STW transition using the identical events as those in the STS transition. For STW, the walking phase started at gait initiation – denoted as the toe-off of the swing-phase (ipsilateral) foot – and ended at the toe-off of the stance-phase (contralateral) foot (**Figure 1D**).

**Figure 1.**
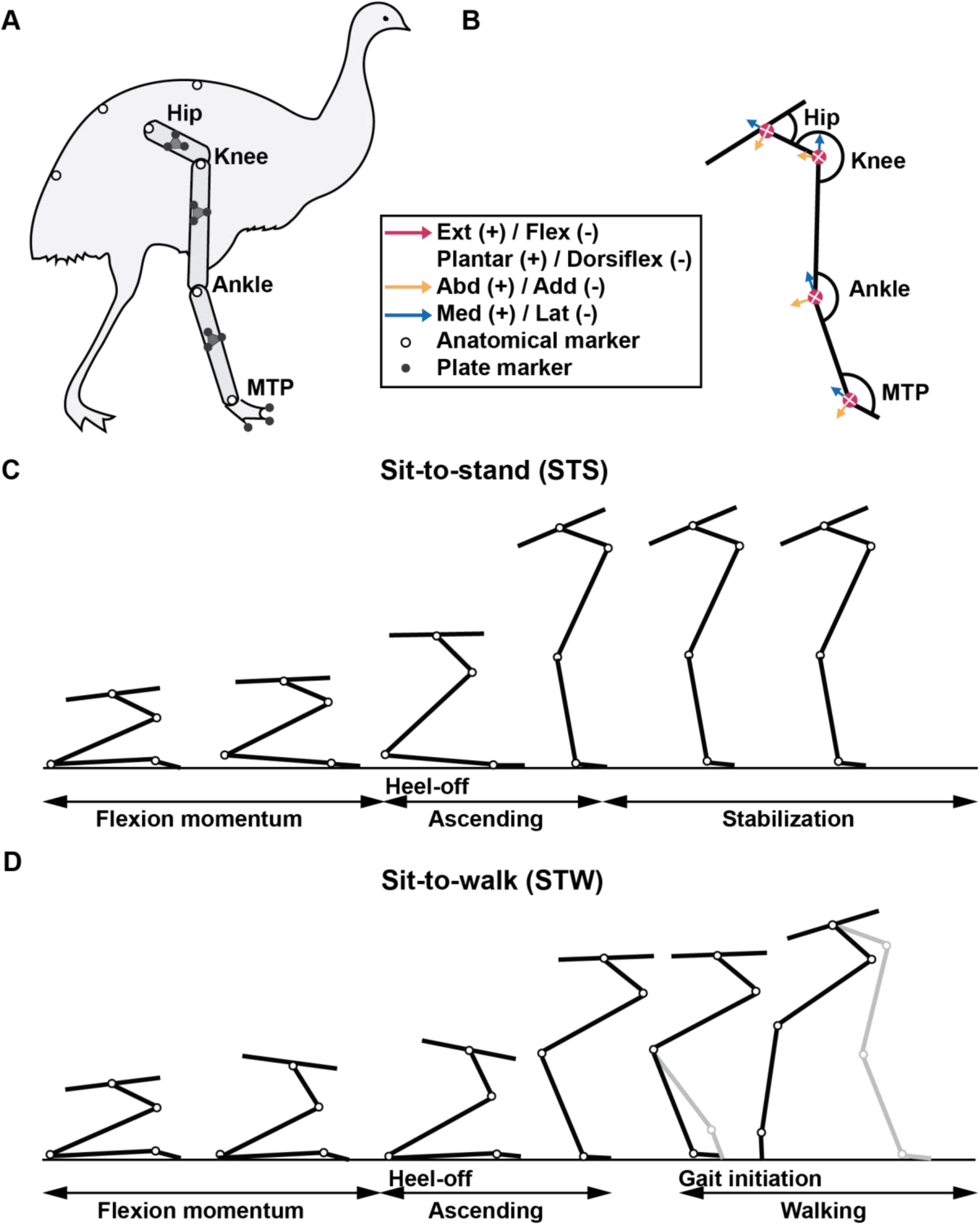
Schematic of (A) emu hindlimb marker locations, (B) joint axes definitions (extension/flexion, plantar/dorsiflexion, abduction/adduction, and lateral [external] or medial [internal] rotation), and (C) STS and (D) STW cycles. The foot position affects how close the COM is to the back edge of the third tarsometatarsophalangeal (MTP) joint. Heel-off in STS and STW are denoted. Gait initiation in STW occurs at the end of the ascending phase and is denoted by the toe-off of the swing-phase foot.

Coordinate data collected from retro-reflective marker clusters placed on the pelvis, right-side femur, tibiotarsus, tarsometatarsus, and digits were used to compute three-dimensional (3D) segment and joint kinematics (**Figure 1A**; marker set definitions in **Supplementary Text**). Marker data were recorded at 250 Hz using 10 Oqus cameras (Qualisys Motion Capture Systems v 2.6.673, Gothenburg, Sweden; ± 1 mm precision) and subsequently filtered (6 Hz fourth-order zero-lag low-pass Butterworth filter) to remove noise. In instances where foot markers were initially obscured, we imputed missing data with the first valid coordinate value. Using OpenSim’s Inverse Kinematics routine (Delp *et al*., 2007), we computed joint kinematics using global optimisation, ensuring that maximum marker errors for bony landmarks were less than 2 cm and root mean square (RMS) errors were less than 1 cm (Hicks *et al*., 2015). Angles were initially represented as Cardan angles of rotation order XY’Z’’ relative to a neutral pose set at 0° (limb fully straightened). These angles were thus measured as positive or negative values relative to this pose and subsequently converted to the convention depicted in (**Figure 1B**).

We recorded GRFs and centre of pressures (COPs) using two Kistler force plates (Model 9287BA, Kistler Holding AG, Winterthur, Switzerland). The force data were initially collected at 500 Hz and then downsampled to 250 Hz for synchronisation with marker data. Subsequently, we conducted baseline removal, and applied a filtering process using a fourth-order zero-lag low-pass Butterworth filter set at 6 Hz to refine the force data. For each trial, both emu hindlimbs were positioned on a single force plate, allowing us to measure the combined hindlimb GRF, free moment and whole-body COP data throughout the movement. Our pilot studies indicated that the emus applied nearly vertical GRFs with each limb pair while standing up, with minimal medio-lateral (ML) GRFs (**Figure S4**). To obtain single hindlimb kinetic data (GRFs, free moments and COPs) for our simulations, we assumed bilateral symmetry (i.e., half of the measured GRFs and free moments were applied to each limb). Next, we calculated the cranio-caudal portion of the COP movement (Supplementary Figure S2) and the ML component of the third digit marker movement to create a composite, single hindlimb GRF dataset following Ellis, Rankin and Hutchinson (2018), ensuring that the right-limb COP remained within the base of support (BOS) consisting of the right tarsometatarsus and digit segments.. Finally, to apply the forces to the two segments forming the foot in the musculoskeletal model, we further partitioned the GRFs and free moments between the two segments to obtain the final GRF motion file (see **Supplementary Text** for details). All data were processed in MATLAB (v2023b, MathWorks, Natick, USA).

We examined only trials where emus began in a near-symmetrical, crouched posture with both tarsometatarsus and digits on the force plate following (Ellis *et al*., 2018), resulting in a potential pool of 3 STS and 9 STW trials (see **Supplementary Text** for detailed inclusion and exclusion criteria). Simulations were generated using each of these trials. For the exemplar STS and STW trials, qualitative assessments were made based on natural movement and data within the observed kinematic and kinetic ranges (**Table 1, Videos S3, Figures S1 and S2**).

**Table 1.**
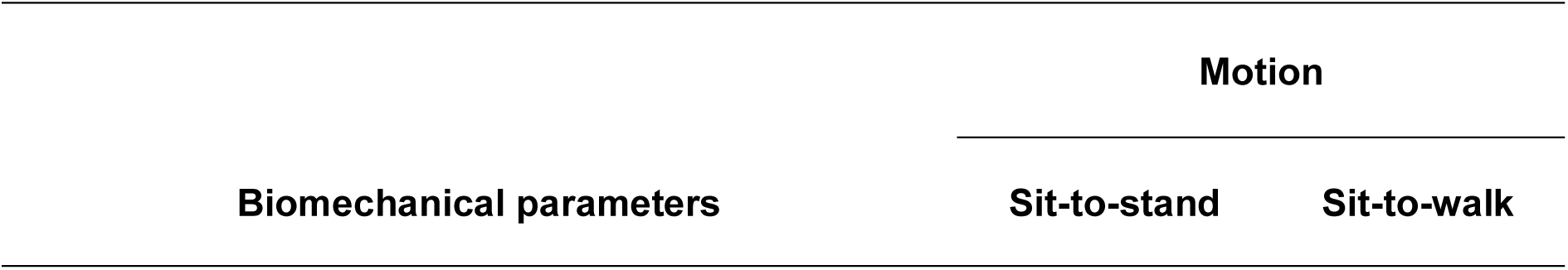
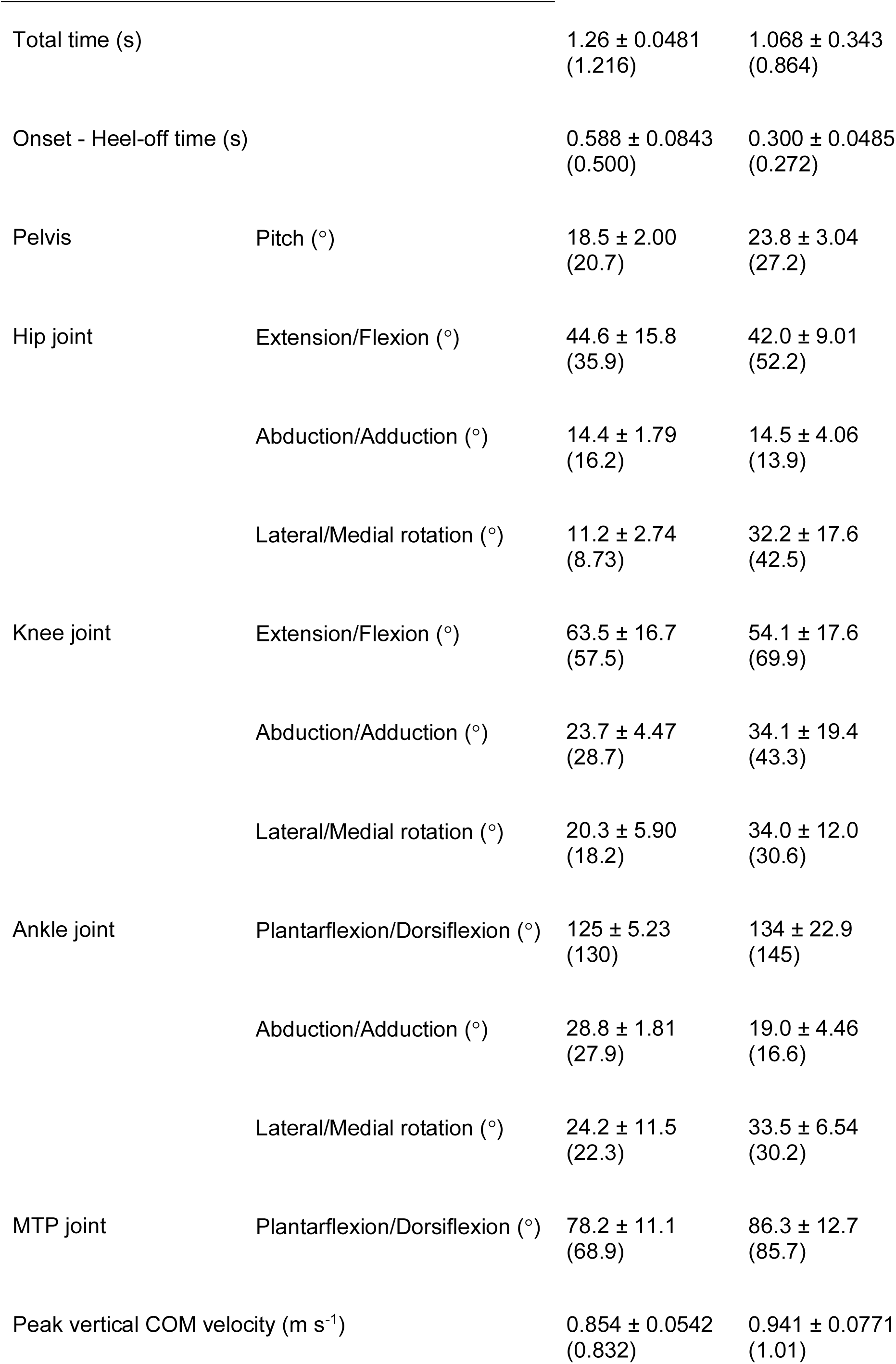
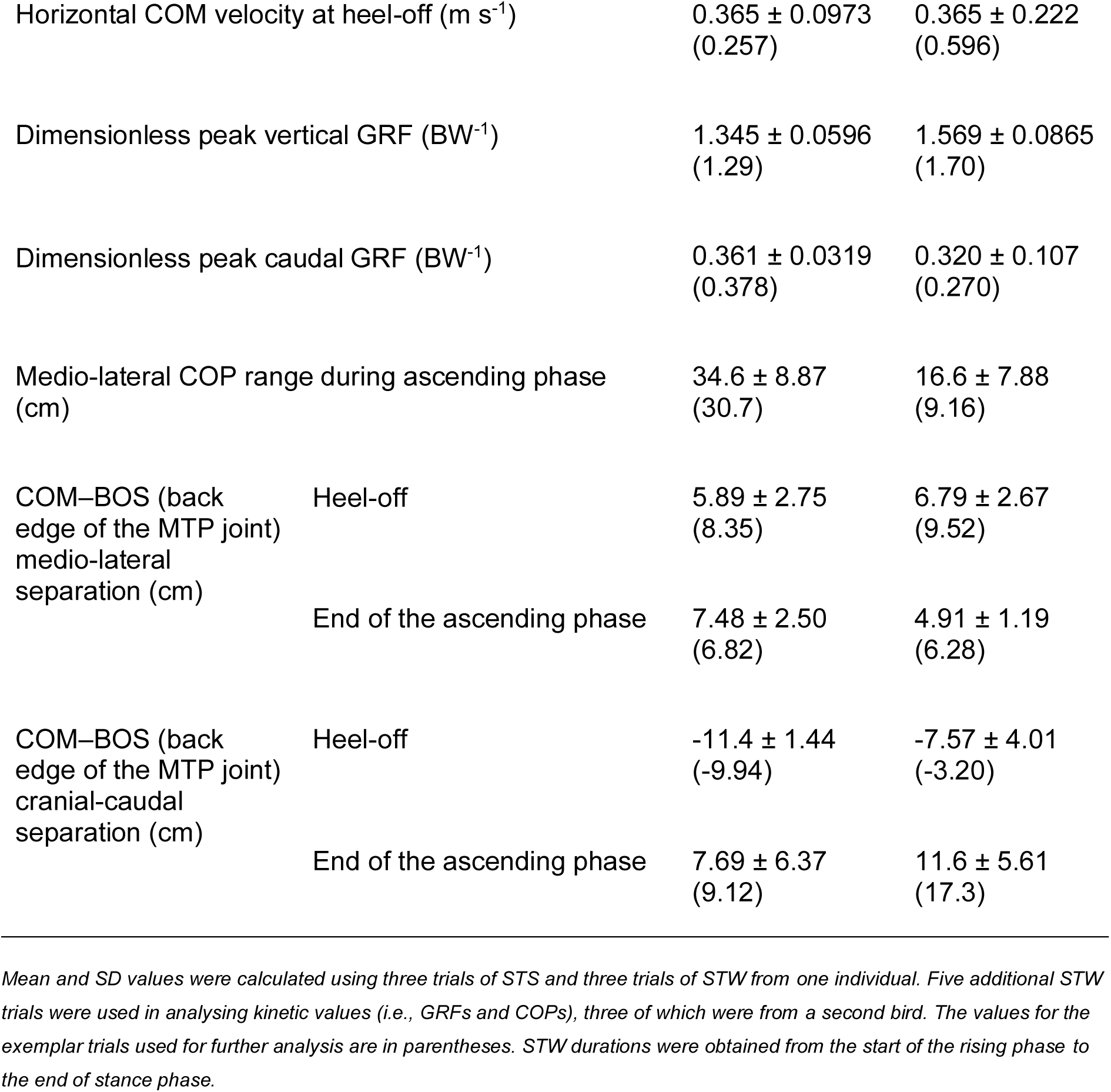
Kinematic and kinetic parameters during the STS and STW transitions.

| Biomechanical parameters | Motion |  |
| --- | --- | --- |
|  | Sit-to-stand | Sit-to-walk |

|  |  |  |  |
| --- | --- | --- | --- |
| Total time (s) |  | 1.26 ± 0.0481<br>(1.216) | 1.068 ± 0.343<br>(0.864) |
| Onset - Heel-off time (s) |  | 0.588 ± 0.0843<br>(0.500) | 0.300 ± 0.0485<br>(0.272) |
| Pelvis | Pitch (°) | 18.5 ± 2.00<br>(20.7) | 23.8 ± 3.04<br>(27.2) |
| Hip joint | Extension/Flexion (°) | 44.6 ± 15.8<br>(35.9) | 42.0 ± 9.01<br>(52.2) |
|  | Abduction/Adduction (°) | 14.4 ± 1.79<br>(16.2) | 14.5 ± 4.06<br>(13.9) |
|  | Lateral/Medial rotation (°) | 11.2 ± 2.74<br>(8.73) | 32.2 ± 17.6<br>(42.5) |
| Knee joint | Extension/Flexion (°) | 63.5 ± 16.7<br>(57.5) | 54.1 ± 17.6<br>(69.9) |
|  | Abduction/Adduction (°) | 23.7 ± 4.47<br>(28.7) | 34.1 ± 19.4<br>(43.3) |
|  | Lateral/Medial rotation (°) | 20.3 ± 5.90<br>(18.2) | 34.0 ± 12.0<br>(30.6) |
| Ankle joint | Plantarflexion/Dorsiflexion (°) | 125 ± 5.23<br>(130) | 134 ± 22.9<br>(145) |
|  | Abduction/Adduction (°) | 28.8 ± 1.81<br>(27.9) | 19.0 ± 4.46<br>(16.6) |
|  | Lateral/Medial rotation (°) | 24.2 ± 11.5<br>(22.3) | 33.5 ± 6.54<br>(30.2) |
| MTP joint | Plantarflexion/Dorsiflexion (°) | 78.2 ± 11.1<br>(68.9) | 86.3 ± 12.7<br>(85.7) |
| Peak vertical COM velocity (m s<sup>-1</sup>) |  | 0.854 ± 0.0542<br>(0.832) | 0.941 ± 0.0771<br>(1.01) |
| Horizontal COM velocity at heel-off ($\text{m s}^{-1}$) | | $0.365 \pm 0.0973$<br>(0.257) | $0.365 \pm 0.222$<br>(0.596) |
| Dimensionless peak vertical GRF ($\text{BW}^{-1}$) | | $1.345 \pm 0.0596$<br>(1.29) | $1.569 \pm 0.0865$<br>(1.70) |
| Dimensionless peak caudal GRF ($\text{BW}^{-1}$) | | $0.361 \pm 0.0319$<br>(0.378) | $0.320 \pm 0.107$<br>(0.270) |
| Medio-lateral COP range during ascending phase (cm) | | $34.6 \pm 8.87$<br>(30.7) | $16.6 \pm 7.88$<br>(9.16) |
| COM–BOS (back edge of the MTP joint) medio-lateral separation (cm) | Heel-off | $5.89 \pm 2.75$<br>(8.35) | $6.79 \pm 2.67$<br>(9.52) |
| | End of the ascending phase | $7.48 \pm 2.50$<br>(6.82) | $4.91 \pm 1.19$<br>(6.28) |
| COM–BOS (back edge of the MTP joint) cranial-caudal separation (cm) | Heel-off | $-11.4 \pm 1.44$<br>(-9.94) | $-7.57 \pm 4.01$<br>(-3.20) |
| | End of the ascending phase | $7.69 \pm 6.37$<br>(9.12) | $11.6 \pm 5.61$<br>(17.3) |

Mean and SD values were calculated using three trials of STS and three trials of STW from one individual. Five additional STW trials were used in analysing kinetic values (i.e., GRFs and COPs), three of which were from a second bird. The values for the exemplar trials used for further analysis are in parentheses. STW durations were obtained from the start of the rising phase to the end of stance phase.

### Musculoskeletal model

We constructed the musculoskeletal model by integrating muscle and tendon architecture, digitised muscle paths, and computed tomography (CT) scan data obtained through dissection (Lamas, 2015). The model comprised five rigid body segments representing the right-side femur, tibiotarsus, tarsometatarsus, and digits as well as the pelvis and remainder of the body (left hindlimb omitted and body mass properties halved, assuming symmetrical support). The model had 10 degrees of freedom (DOF) representing the right hip (3 DOF), knee (3 DOF), ankle (3 DOF), and tarsometatarsophalangeal (MTP) (1 DOF) joints. In the model, the pelvis moved freely relative to the ground (three rotational and three transitional DOFs); however, we constrained the pelvis’s roll movement during simulations, assuming bilateral symmetry. We adjusted segment mass, inertia, length, and musculotendon length values by scaling the original model in OpenSim to match the body masses and bone lengths of the bird used during experimental data collection (see **Supplementary Text**).

Our emu hindlimb musculoskeletal model encompassed 37 muscle-tendon unit (MTU) actuators and 19 MTU wrapping objects representing 33 muscles (**Figure 2**, **Table S1**). Each MTU actuator was represented using a Hill-type model with intrinsic force-length-velocity relationships (Millard *et al*., 2013), with muscle fascicle lengths (assumed to equal optimal fibre length), masses, and pennation angles measured through dissection of the same individual used for model construction, except for M. ischiofemoralis (ISF), where parameters were sourced from another individual of the same body mass.

**Figure 2.**
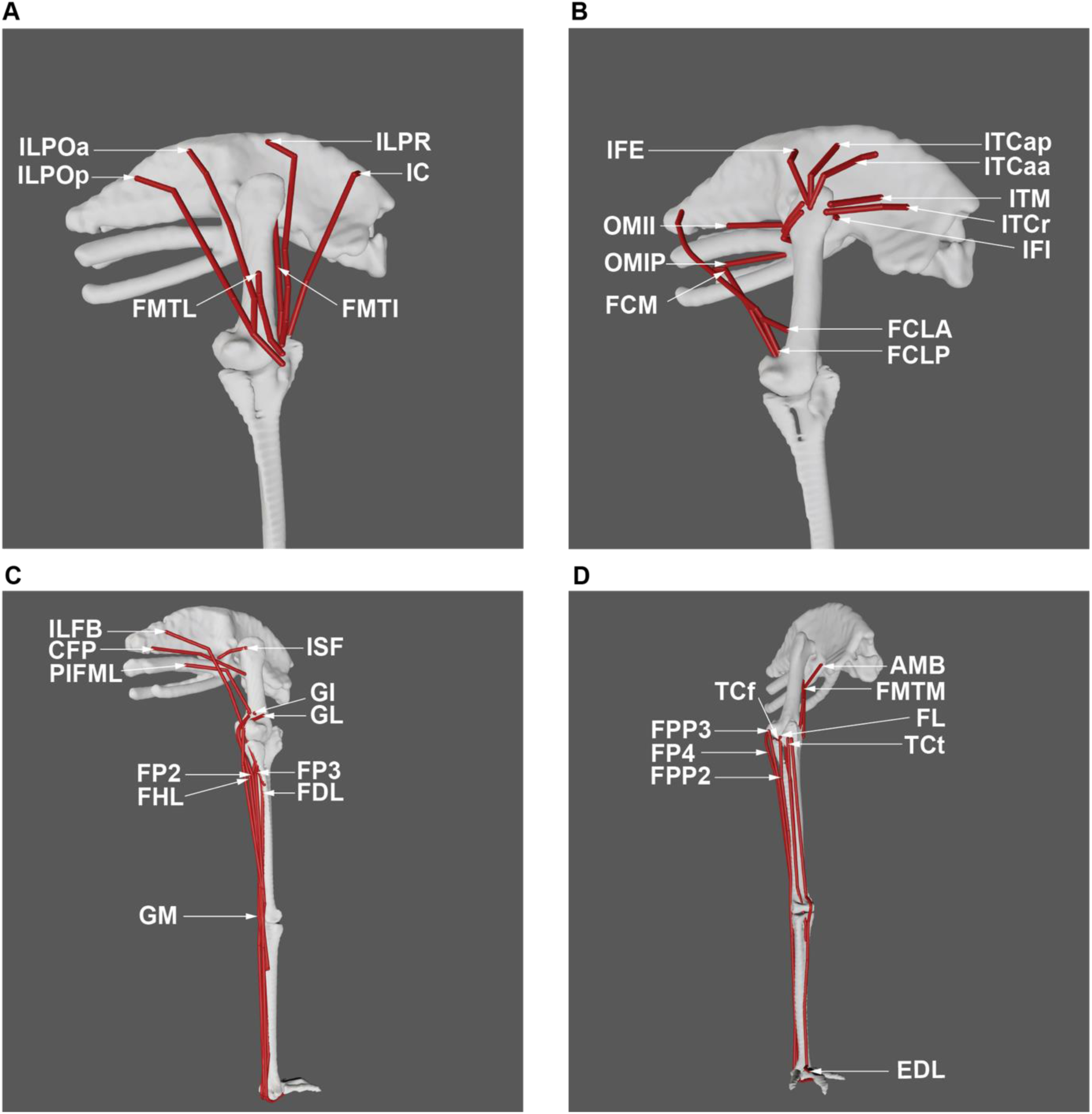
Muscle groups included in the musculoskeletal model of the right hindlimb of the representative emu, in the neutral pose with initial muscle attachments used. See **List of symbols and abbreviations** and **Table S1** for muscle abbreviations. (A) ‘Triceps femoris’ knee extensor muscles (except AMB, FMTM), in lateral view. (B) Deep dorsal (IFE, ITCaa, ITCap, ITCr, ITM, IFI), OMII, OMIP, and ‘hamstring’ (FCM, FCLP, FCLA) thigh muscles, in craniolateral view. (C) Other caudally positioned pelvic muscles, and distal hindlimb muscles on the plantar (caudal) surface of the limb, in caudolateral view. (D) AMB and distal hindlimb muscles on the dorsal (cranial) surface of the limb, in craniolateral view.

Following assignment of architectural properties and definition of MTU paths, we estimated TSLs for each MTU across diverse behaviours (e.g., walking, sit-to-stand, and sit-to-walk) based on (Manal and Buchanan, 2004). TSL is an abstraction of the resting length of the modelled tendon, at which zero force is generated. In some cases this approach estimated a negative value for M. iliofemoralis internus (IFI): TSL was then recalculated using 5 % of the static trial MTU length as an initial value. We fine-tuned TSLs with the goal of having normalised fascicle lengths operate within the range of 0.5 – 1.5 (Hicks *et al*., 2015) throughout each simulation and be between 0.8 and 1.2 in the final posture (Ellis *et al*., 2018). Twenty-five of the 37 muscles were adjusted, with an average TSL adjustment of 5.2 % (SD 6.6 %) of the originally estimated MTU length (**Table S1**). For some muscles, muscle fibre lengths remained unrealistic throughout the motion (i.e., < 0.5 or > 1.5 times resting length) even after adjustments to TSL. In these instances, adjustments were made to optimal fibre length, concurrently shortening or lengthening TSL to maintain a constant MTU length, to ensure that muscles operated between 0.5 and 1.5 times their optimal length. This fibre length adjustment was done for M. flexor cruris lateralis pars accessoria (FCLA) (10 % increase), M. flexor cruris medialis (FCM) (50 % increase), M. iliofemoralis externus (IFE) (80 % increase), M. puboischiofemoralis p. lateralis and p. medialis (PIFML) (17 % increase), M. flexor perforans et perforatus digiti III (FPP3) (2 % increase), and M. flexor perforatus digiti IV (FP4) (70 % increase) muscles. Each muscle’s maximal isometric force (*F_max_*) was calculated according to the formula:

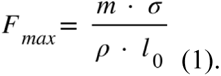

where *m* denotes muscle belly mass, *σ* represents maximum isometric stress of the fibres and *ρ* is muscle tissue density. Values of *σ* (300,000 N m^-2^; Medler, 2002; Hutchinson, 2004) and *ρ* (1,060 kg m^-3^; Mendez and Keys, 1960; Hutchinson *et al*., 2015) that are standard for vertebrate skeletal muscle were used. Note that pennation angle is not used in Eqn 1 because OpenSim incorporates it as a separate parameter.

We incorporated three additional actuators to compensate for the residual forces and moments at the pelvis and four reserve actuators – one for each DOF in the right limb – to compensate for mechanical work that could not be solely satisfied by the muscles (Hicks *et al*., 2015). Our optimisation framework focused on minimising the use of these ‘reserve actuators’ to ensure that required joint moments were predominantly supplied via muscles.

### Simulations

We simulated four trials (2 STS, 2 STW) using the experimental exemplar STS and STW data and musculoskeletal model. For each trial two different optimisation frameworks were used to estimate MTU activations, force, and length changes (detailed below), resulting in 8 simulations in total. Further sensitivity analyses involving additional simulations are presented later.

In the first optimisation framework, we used the static optimisation routine in OpenSim (Delp *et al*., 2007) to generate each simulation. This method optimised a pre-set objective criterion, minimising the sum of squared muscle activations across all muscles at each time step while adhering to biomechanical constraints including static equilibrium (Delp *et al*., 2007). With a time step set to 0.005 s, the simulations provided MTU activation, force, and length time histories throughout the movement cycle. This approach worked independently at each time step, excluding energy transfer between steps (e.g., tendon energy storage and return) and muscle excitation-activation dynamics. Additionally, this framework disregarded passive fibre force generation and assumed rigid tendons, resulting in muscle fibre length changes needing to account for all MTU length changes.

In the second optimisation framework, we performed dynamic simulations (*sensu* De Groote *et al*., 2016; Peter J. Bishop *et al*., 2021) using OpenSim Moco (Dembia *et al*., 2020). Using the same objective criterion as the static optimisation routine, this approach factored in the model state from prior time steps (e.g. joint angles, muscle activation level, tendon strain) to influence the optimal solution for the current step, similar to purely forward dynamic simulations. Linking time steps allowed for the integration of muscle excitation – activation dynamics, consideration of non-rigid tendon characteristics, and incorporation of passive muscle fibre force generation, which was a more realistic approach.

### Analysis

In analysing kinematic and kinetic parameters during STS and STW, we computed mean and standard deviation (SD) values using three trials of STS and three trials of STW from one individual (due to limited data, see **Supplementary Text**). For kinetic values (i.e., GRFs and COPs), five more STW trials were included, three of which were from a second bird. For STS trials, we calculated range of motion (ROM) values from the start of the movement to the end of the stabilisation phase. For STW trials, we calculated ROM values for the stance limb, from the start of the movement to stance toe-off.

We performed a series of sensitivity analyses to test our main modelling and simulation assumptions. The static optimisation simulations assumed that tendons were inextensible and muscles did not generate passive forces. To test the robustness of this assumption, we varied TSLs by ± 5 % following (Scovil and Ronsky, 2006; Redl, Gfoehler and Pandy, 2007; also Ellis, Rankin and Hutchinson, 2018), and muscle dynamics were compared across conditions to assess how TSL changes altered them. To reduce the potentially confounding factors of different tendon and muscle models when directly comparing between static optimisation and dynamic (i.e., the direct collocation in Moco) simulations, we generated the following additional simulations. These simulations were identical to the previous dynamic simulations but introduced variations: one had limited tissue properties which eliminated tendon-muscle fibre dynamics and partially negated the ability of a forward dynamics optimisation to account for time-dependent muscle interactions (e.g. tendon energy storage and return), and the other incorporated non-rigid tendons but eliminated passive muscle fibre force generation capability. As a result, these simulations would not be a realistic choice under normal circumstances. However, being able to control for these variables allowed us to eliminate these potentially confounding factors to make a more direct comparison between the static optimisation and inverse simulation frameworks. Indeed, this controlled approach allows us to gain additional insight into the roles of the different structures of the musculotendon system. Finally, we performed a nominal static simulation on two additional STS/STW trials from the same emu.

In evaluating the potential impact of reserve actuators on simulation outcomes, we compared reserve actuator values with the net joint moments derived from OpenSim’s inverse dynamics analysis. This comparison involved calculating average reserve actuator values and net joint moments and examining reserve actuator values at the instances of peak net joint moments in each DOF.

## Results

### Kinematics

Analysis of the experimental kinematic data revealed similarities and noticeable differences in the STS and STW movement trajectories (**Table 1**, **Figure 3**). Both movements involved large excursion ranges at each joint. Extension in the hip and knee joints occurred during the early stage of the transition, reaching their peaks near heel-off, followed by a small flexion and subsequent extension towards the later stage of the movements. The ankle joint exhibited the most substantial excursion among all joints, reaching 125 ± 5.23° (mean ± 1 SD) in STS and 134 ± 22.9° (mean ± 1 SD) in STW, extending predominantly from heel-off throughout the remainder of each motion. The MTP joint initially maintained a stationary plantarflexed posture, and then transitioned into dorsiflexion at the time of heel-off. STW was a fluid merging of STS and gait. At gait initiation, the hip, knee, and ankle remained flexed in STW compared to the upright standing pose of STS, and emus exhibited larger hip and knee joint non-sagittal motions during the STW transition to walking, although non-sagittal motions were relatively small compared to sagittal motions in both movements. In STW, a rapid flexion in the hip and knee joints, dorsiflexion in the ankle joint, and plantarflexion in the MTP joint occurred right before the stance-phase limb’s toe-off. STW also exhibited a shorter flexion momentum period (i.e., from onset of the movement to heel-off) (STS: 0.588 ± 0.0843 s; STW: 0.300 ± 0.0485 s; mean ± 1 SD), larger peak COM velocity (STS: 0.854 ± 0.054 m s-1; STW: 0.941 ± 0.0771 m s-1; mean ± 1 SD), and a noticeable body downward pitch at the onset of the movement. Furthermore, simulations results indicated that the COM– BOS separation (i.e., distance between the whole-body COM and the back edge of the third tarsometatarsophalangeal joint) was smaller in the cranial-caudal direction and larger in the medio-lateral direction at heel-off in STW (cranial-caudal direction: −7.57 ± 4.01 cm; medio-lateral direction: 6.79 ± 2.67 cm; mean ± 1 SD) compared to STS (cranial-caudal direction: - 11.4 ± 1.44 cm; medio-lateral direction: 5.89 ± 2.75 cm; mean ± 1 SD).

**Figure 3.**
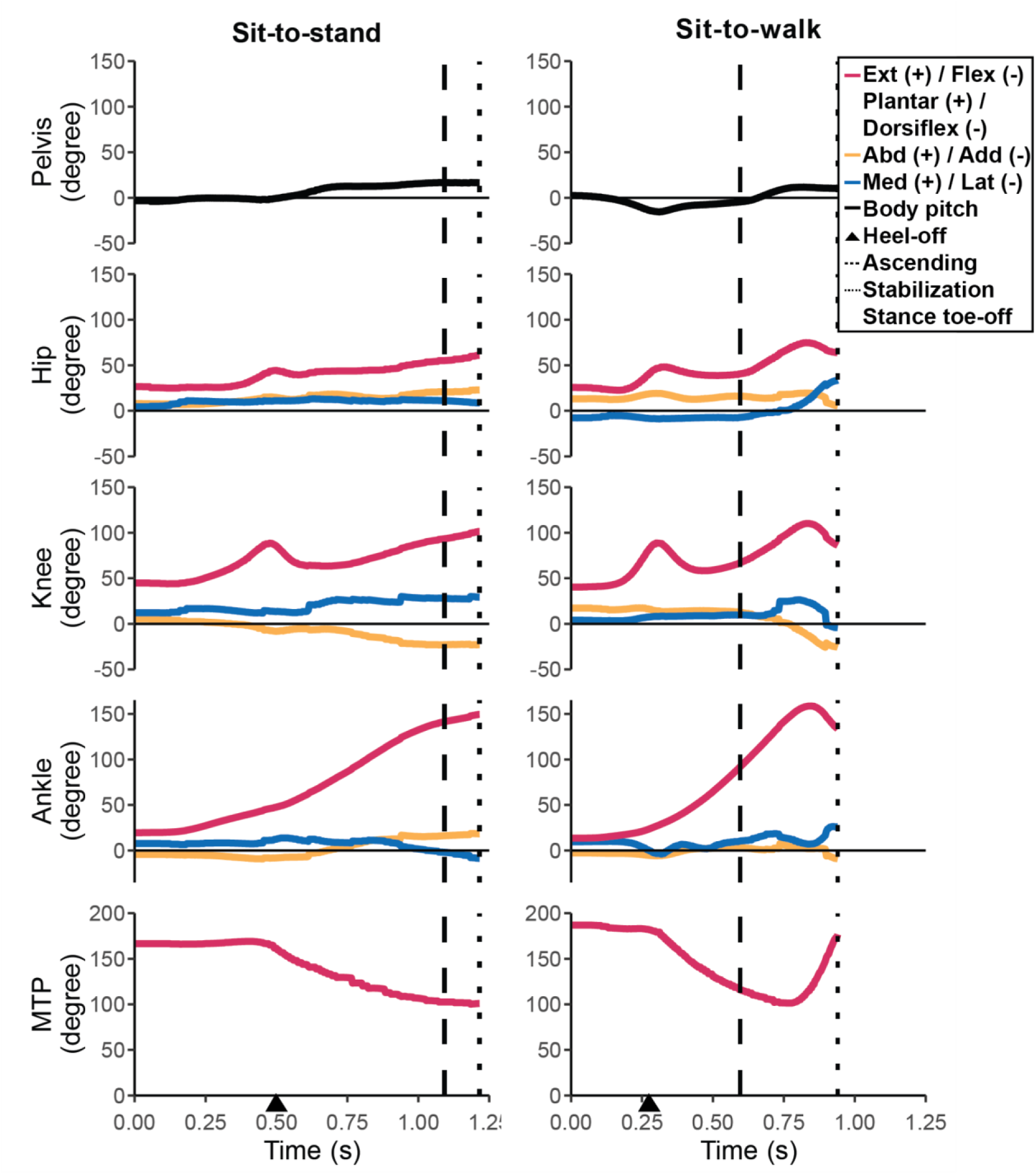
Body and hindlimb joint angles during the STS and STW transitions in the exemplar trials. See Figure 1 for marker/angle definitions. For STW, the results were derived from the stance leg. The STS and STW events and phases are denoted, where heel-off is represented by an arrow, the end of the ascending phase is represented by a dashed line, and the ends of the stabilisation phase of STS and walking phase of STW are represented by dotted lines.

### Kinetics

In all directions, the GRFs reached a peak early in the movements, occurring around heel-off (**Table 1**, **Figure 4**). Emus initiated forward momentum with a moderate cranially oriented GRF, while experiencing relatively small medio-lateral GRF throughout both movements. STW displayed a larger peak vertical GRF compared to STS, nearly approaching twice the body weight (BW) (STS: 1.345 ± 0.0596 BW; STW: 1.569 ± 0.0865 BW; mean ± 1 SD), comparable to values reported for walking and slower speed running (Goetz *et al*., 2008), and a smaller braking impulse (STS: 0.361 ± 0.0319 BW; STW: 0.320 ± 0.107 BW; mean ± 1 SD). STW also displayed a smaller medio-lateral COP (STS: 34.6 ± 8.87 cm; STW: 16.6 ± 7.88 cm; mean ± 1 SD). Additionally, a second, smaller vertical GRF peak (∼ 1.3 BW) occurred at the onset of STS, potentially attributed to the initial adjustment of foot position closer to the COM in the exemplar trials. Despite this anomaly, we considered the exemplar trial suitable for subsequent simulations due to its natural kinematics and kinetics.

**Figure 4.**
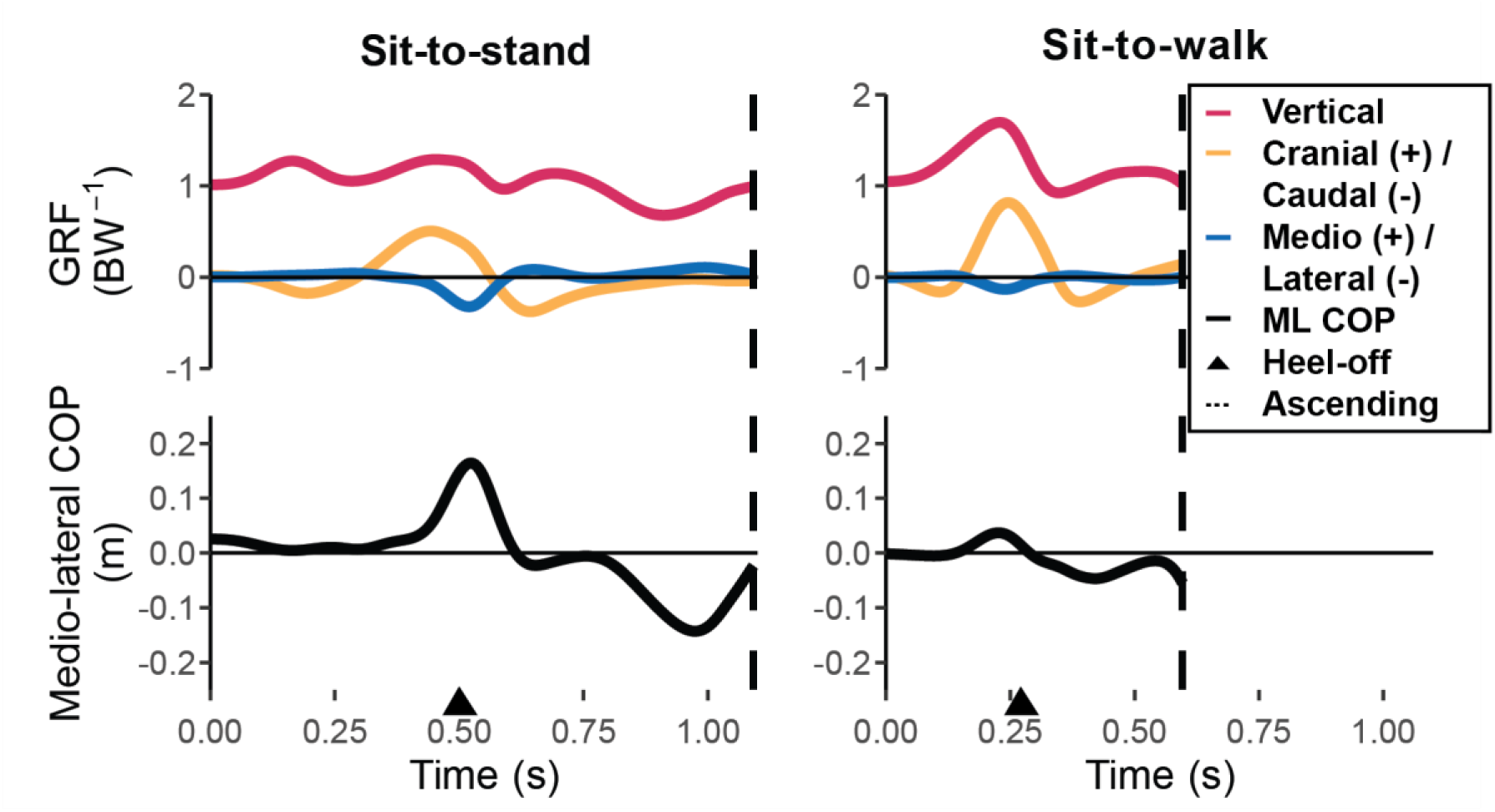
Total (dimensionless) ground reaction forces and medio-lateral (ML) COP during the STS and STW transitions in the exemplar trials. The STS and STW events and phases are denoted, where heel-off is represented by an arrow and the end of the ascending phase is represented by a dashed line.

Consistent with the GRF profile, our inverse dynamics analysis indicated the need for large net extensor moments at the hip and ankle joints during the ascending phase, especially around heel-off (**Figure 5**). The hip joint exhibited a peak extensor moment (∼ 0.14 dimensionless unit in STS and ∼ 0.21 dimensionless unit in STW) immediately before heel-off, sharply decreasing to near zero, then maintaining a small extensor moment (< 0.1 dimensionless unit) during the later ascending phase. The ankle joint sustained a relatively large extensor moment throughout STS and STW, peaking at ∼ 0.18 dimensionless unit in STS and ∼ 0.27 dimensionless unit in STW right after heel-off. In contrast, the knee joint required a relatively small extensor moment (< 0.1 dimensionless unit) and exhibited a peak flexor moment (< 0.1 dimensionless unit in STS and ∼ 0.14 dimensionless unit in STW) around heel-off. The MTP joint required a minimal plantarflexor moment (< 0.1 dimensionless unit). With the exception of the hip joint lateral rotator and adductor moments, exemplar trials revealed generally modest non-sagittal moments in both movements.

**Figure 5.**
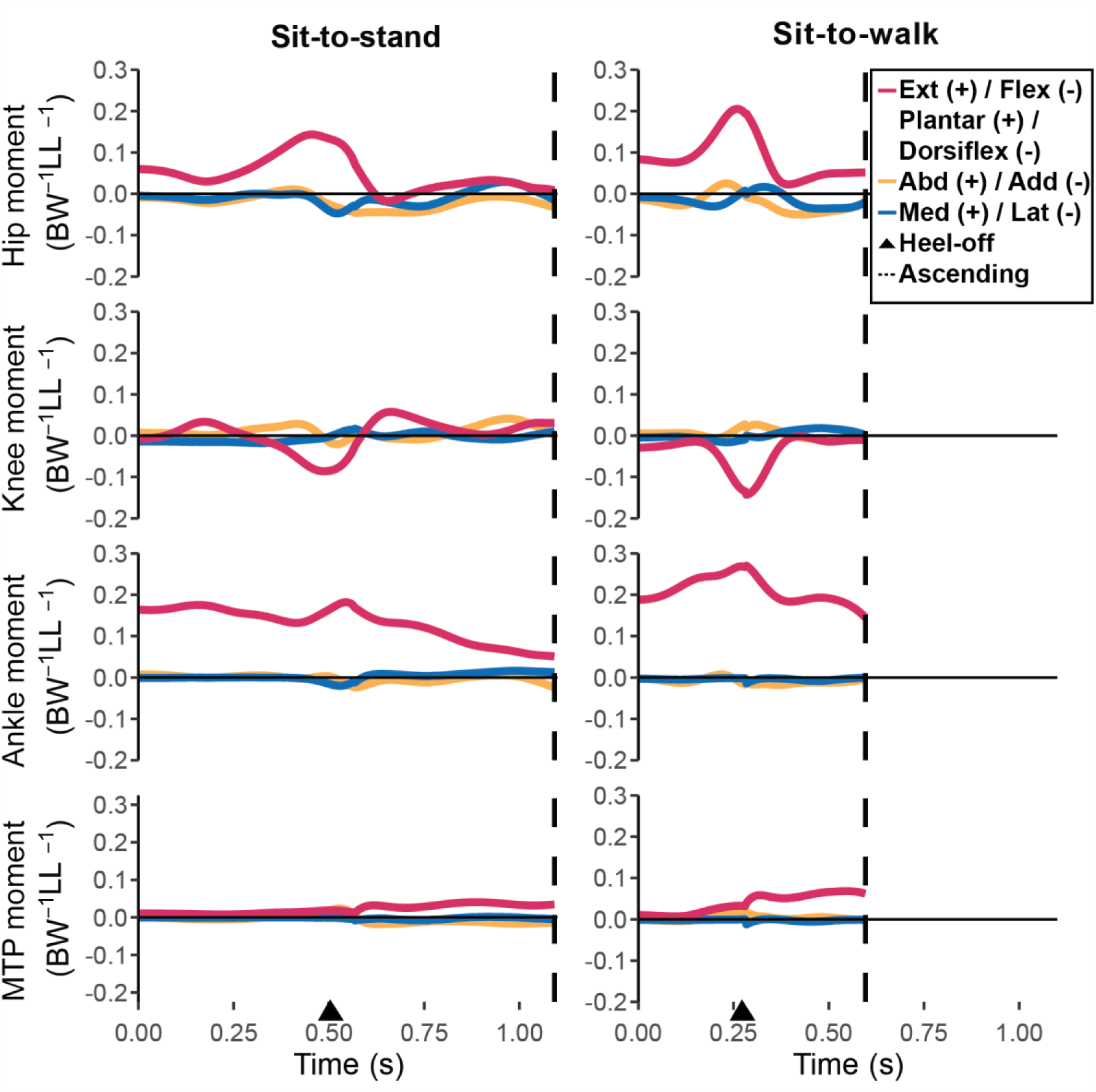
Net (dimensionless) joint moments during the STS and STW transitions in the exemplar trials. For STW, the results were derived from the stance leg. The STS and STW events and phases are denoted, where heel-off is represented by an arrow and the end of the ascending phase is represented by a dashed line.

### Muscle activations, length changes, and forces

Simulated muscle activation patterns were broadly similar across the different simulation frameworks. Consistent with the joint moment pattern (**Figure 5**), most simulations estimated muscle activations and forces peaked around heel-off, then sharply decreased. A few muscles remained active through the end of the movements (**Figures 6 – 9**, **Figure S3**). We focus here on results of the nominal static optimisation simulation and dynamic simulation with full tissue properties and discuss sensitivity analyses later.

**Figure 6.**
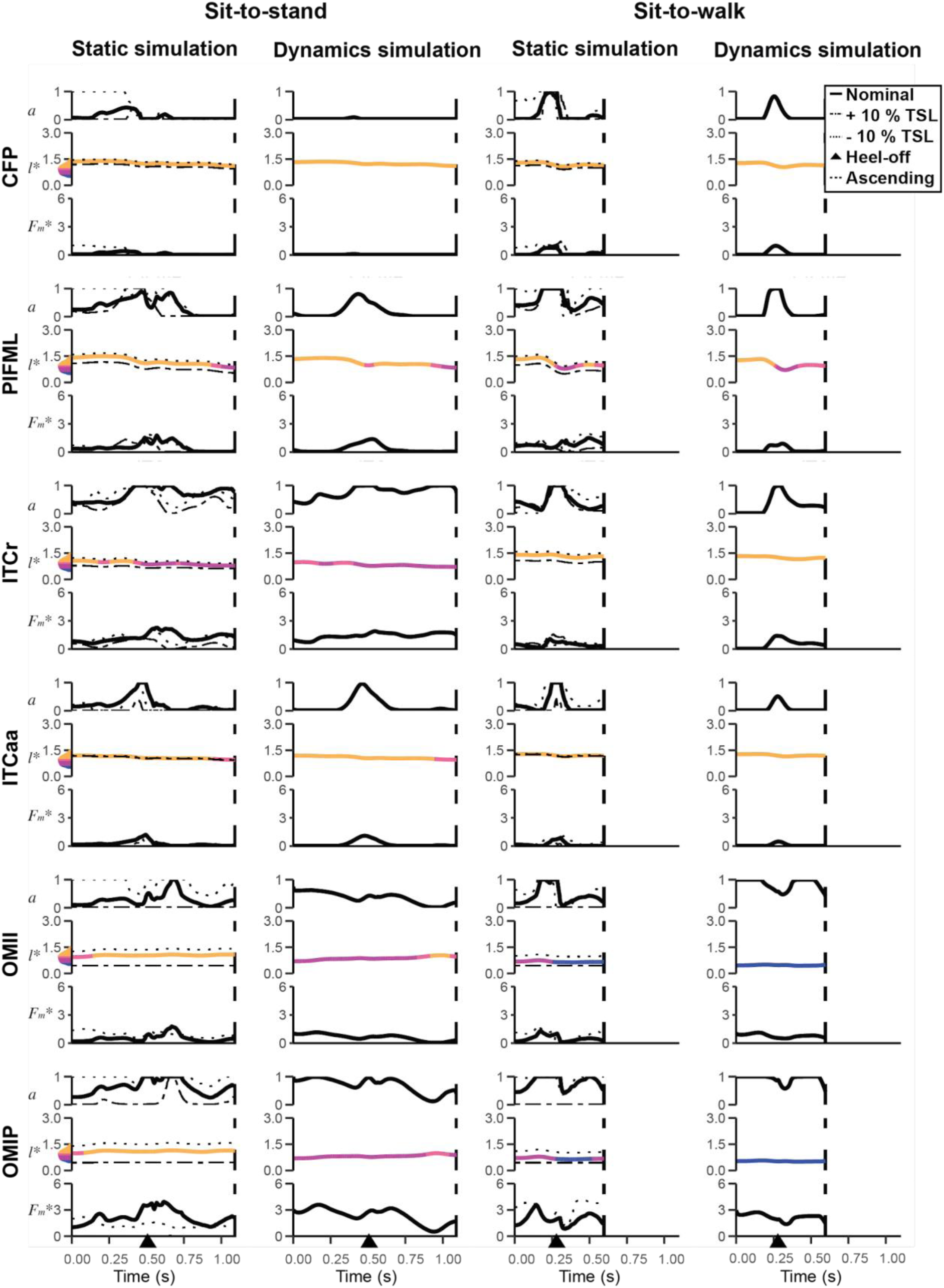
Simulated muscle activations (*a*), normalised fibre lengths (*l\**), and body weight-nondimensionalised muscle forces (*F_m_\**) of uniarticular hip muscles during the STS and STW transitions in the exemplar trials. For STW, the stance leg was simulated. Results from static simulations (solid line: nominal simulation; dashed line: + 5 % TSL; dotted line: - 5 % TSL) and dynamic simulations with full tissue properties are shown. Normalised fibre lengths are colour coded according to where on the active force-length curve fibres are operating: steep ascending limb, shallow ascending limb, plateau, and descending limb (divisions approximately correspond to (Arnold and Delp, 2011)). The STS and STW events and phases are denoted, where heel-off is represented by an arrow and the end of the ascending phase is represented by a dashed line. Only muscles where simulated activation exceeded 50 % of maximum in dynamic simulation, in either STS or STW, are shown. Muscle abbreviations are defined in **List of symbols and abbreviations** and **Table S1**.

In both nominal static and dynamic simulations of STS and STW, seven of 37 extensor muscles reached near maximal activation values (∼ 1.0). These muscles included one uniarticular hip extensor (**Figure 6**): M. puboischiofemoralis p. lateralis and p. medialis (PIFML), one biarticular hip and knee extensor (**Figure 7**): M. iliotibialis lateralis pars preacetabularis (ILPR), three uniarticular knee extensors (**Figure 7**): M. femorotibialis lateralis (FMTL), intermedialis (FMTI) and medialis (FMTM), and two ankle extensors (**Figure 8**): M. gastrocnemius lateralis (GL) and M. gastrocnemius medialis (GM). Intriguingly, some small, short-fibred hip muscles showed brief but near-maximal activations at the onset of both STS and STW (∼ 1.0) (**Figures 6 and 7**): M. iliotrochantericus cranialis (ITCr), M. iliotrochantericus caudalis anterior part (ITCaa), M. ambiens (AMB), M. obturatorius medialis ilium—ischium part (OMII) and ischium—pubis part (OMIP). We also observed co-activations in the antagonist muscles (**Figures 8 and 9**) M. tibialis cranialis c. femorale (TCf), c. tibiale (TCt) (acting mainly as ankle dorsiflexors but also knee extensors) and M. extensor digitorum longus (EDL) (acting as digit dorsiflexors and ankle dorsiflexors). In general, STW exhibited higher muscle activations than STS (except for ILPR, ITCr, ITCaa, and FP3), particularly early in the transition, with M. caudofemoralis p. pelvica (CFP) (acting as hip extensor) and M. flexor perforans et perforatus digiti III (FPP3) (acting as ankle extensor and digit plantarflexor) also reaching maximal possible activations. In particular, M. gastrocnemius medialis (GM) remained near maximum activation throughout the entire STW transition. Both the nominal static and dynamic simulations predicted most other muscles to have lower activations (< 0.5 peak values; **Figure S3**). There was an overall trend for reduced activations in dynamic simulations with full tissue properties compared to static simulations, except for the distal muscles M. fibularis longus (FL), M. tibialis cranialis c. femorale (TCf), and M. tibialis cranialis c. tibialis (TCt).

**Figure 7.**
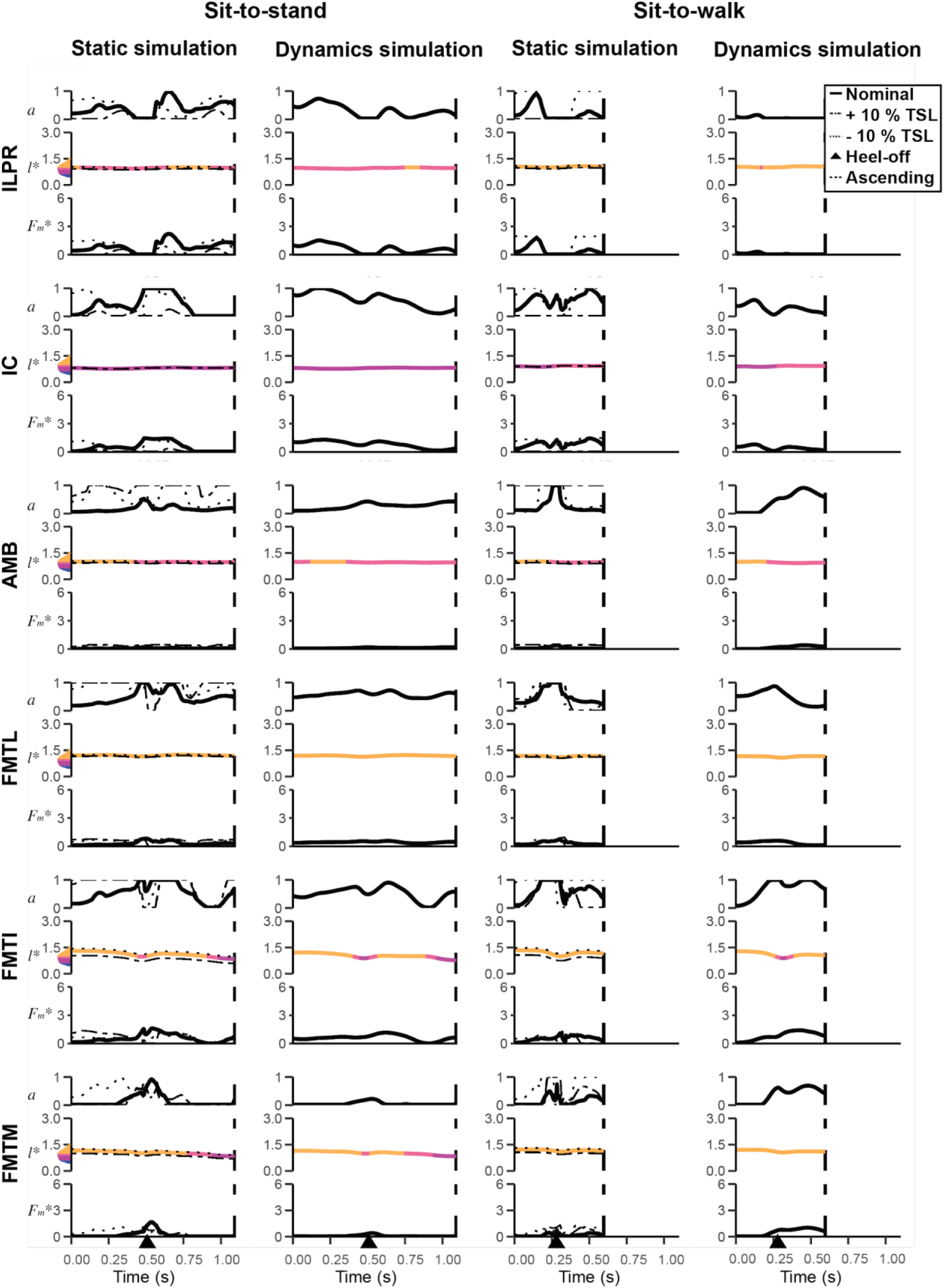
Simulated muscle activation, normalised fibre lengths, and body weight-nondimensionalised muscle force of biarticular muscles crossing the hip; and knee and uniarticular knee muscles; during the STS and STW transitions in the exemplar trials. See Figure 5 for further details.

**Figure 8.**
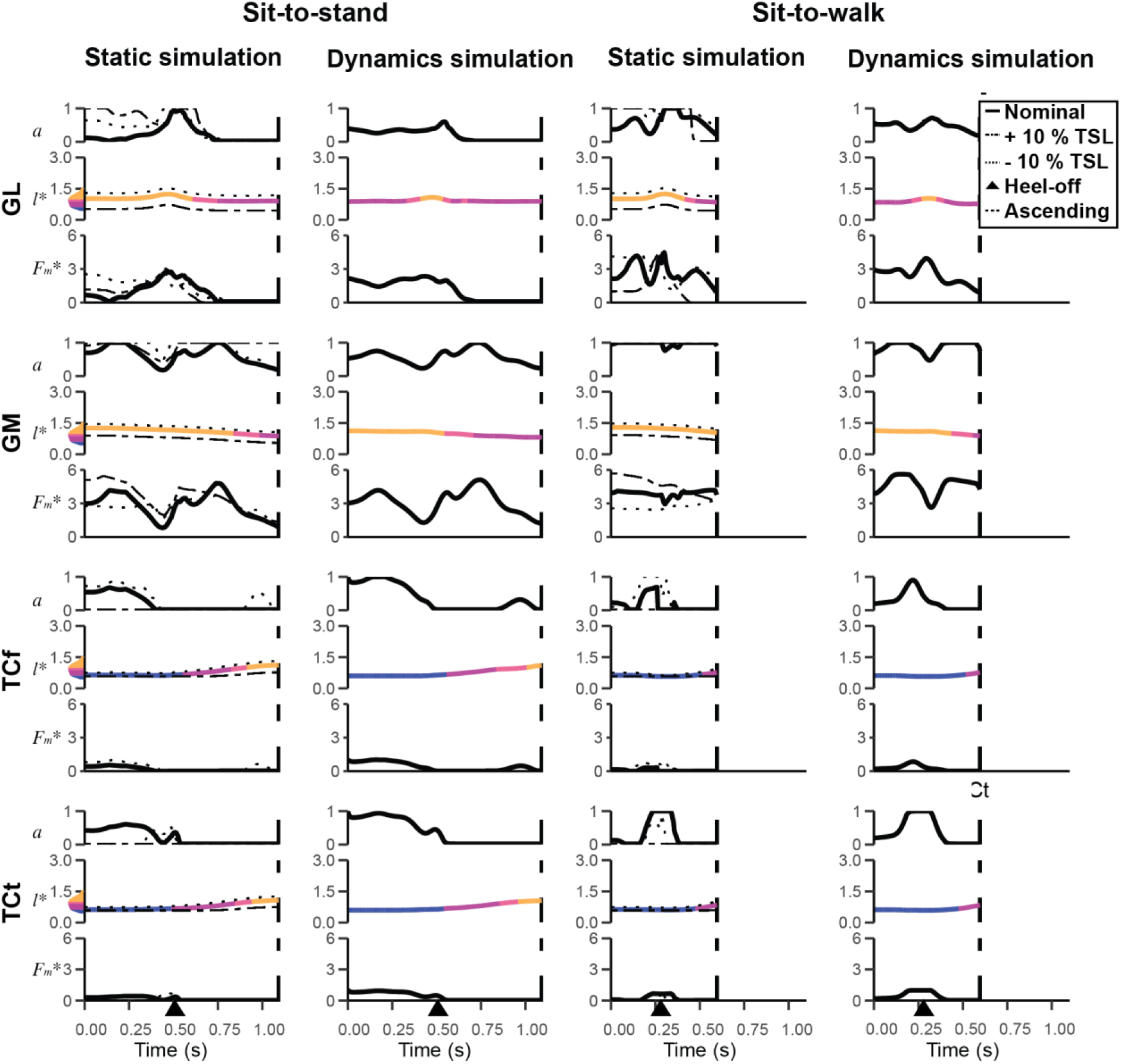
Simulated muscle activation, normalised fibre lengths, and body weight-nondimensionalised muscle force of biarticular muscles crossing the knee and ankle during the STS and STW transitions in the exemplar trials. See Figure 5 for further details.

STS and STW showed substantial changes in muscle fibre lengths in both nominal static and dynamic simulations (**Figures 6 – 9**). All muscle fibres operated within the range of 0.5 – 1.5 times optimal fibre length (as they were tuned to do), with some muscles using a greater portion of this range than others, especially distal limb muscles like FPP3, which showed nearly a 50 % change in its muscle fibre length. We also observed unique patterns of lengthening and shortening across muscle groups at distinct stages of the transition. In static simulations of both movements, the hip extensors CFP and PIFML, knee extensor FMTI, ankle extensor GM and digit plantarflexor FPP3 were at extremely long fascicle lengths early in the transition (∼ 1.5 times optimal fibre length); ankle dorsiflexors TCf and TCt operated at their minimum lengths (∼ 0.5 times optimal fibre length); other muscles (e.g., ILPR) remained at lengths closer to optimal for force generation. At the hip and knee joints, the hip extensors CFP and PIFML and knee extensors FMTI and FMTM shortened right before heel-off, lengthened, and then remained shortened by the end of the movements, whereas the knee extensors FMTL and ILPR (ILPR also acted as a hip flexor) remained nearly isometric throughout the transition. At the ankle, GL lengthened right before heel-off, shortened, and then remained static, whereas GM uniformly shortened. At the digits, FL and FPP3 shortened right before heel-off, following a similar pattern to the uniarticular hip/knee extensors, whereas FP3 lengthened right before heel-off, shortened, and then remained lengthened at the end of the movements. Overall, dynamic simulations with full tissue properties had reduced muscle fibre length changes compared to static simulations, especially in distal limb muscles such as GM, which kept fibres operating at more optimal ranges in those muscles.

**Figure 9.**
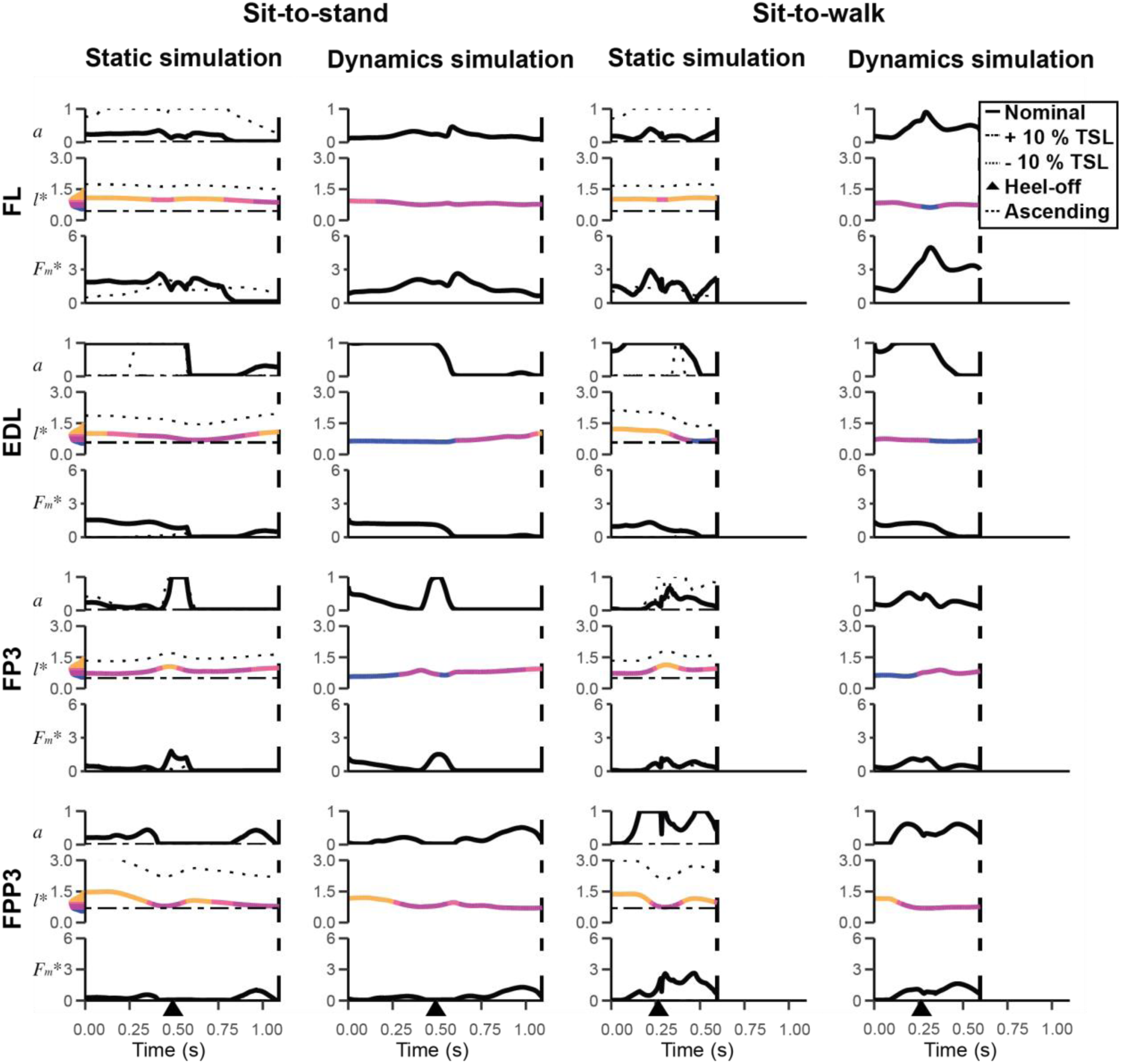
Simulated muscle activation, normalised fibre lengths, and body weight-nondimensionalised muscle force of muscles crossing the MTP joint during the STS and STW transitions in the exemplar trials. See Figure 5 for further details.

In the static simulations, many muscles generated large forces, approaching or exceeding body weight (**Figures 6 – 9**). GM stood out, reaching a maximum force exceeding 4 times body weight, while the major extensor muscles PIFML, ILPR, FMTI, FMTM, GL, FL, and FP3 reached maximal forces of more than 1.5 times body weight. STW generally necessitated larger forces in most muscles compared to STS; FPP3, for example, exerted maximal forces of ∼ 2.5 BW in STW but only ∼ 1.0 BW in STS. The small hip muscles ITCr, ITCaa, OMII, and OMIP also generated moderate (∼ 1.0 BW) to large forces, with OMIP peaking at more than 3.5 times body weight in both STS and STW. AMB had strikingly high activations during STW but only generated small peak forces (∼ 0.4 BW). The largest absolute forces were generated by GM (1342 N in STS and 1170 N in STW), GL (765 N in STS and 1253 N in STW), OMIP (1097 N in STS and 1002 N in STW), and FL (737 N in STS and 821 N in STW), although more than 300 N peak forces were also generated by IC, ILPR, FMTI, OMII, PIFML, and EDL in both movements (and FP3, FMTM and FDL in STS only; ITCaa and ITCr in STW only). Simulations predicted smaller peak forces for CFP (118 N in STS and 281 N in STW), FMTL (227 N in STS and 217 N in STW), TCf (154 N in STS and 89 N in STW), and TCt (122 N in STS and 188 N in STW). Overall, dynamic simulations had reduced muscle forces in most simulated muscles compared to static simulations, with a few exceptions such as GM and FL (**Figures 8 and 9**).

### Reserve actuators

With few exceptions, reserve actuator values in the nominal static and dynamic simulations remained small in comparison to the inverse dynamics joint moments (typically < 1 Nm or ≤ 10 % of average or peak inverse dynamics moments) (**Table S2**). The main exception occurred at the ankle joint, where ankle abduction/adduction reserve actuator moments, comprising 130 % of the total in STS and 57 % in STW, were required in the static optimisation simulation. Similarly, the ankle long-axis rotation reserve actuator accounted for 58 % in STS and 95 % in STW in static optimisation simulation. Generally, altering TSL by ± 5 % increased the demand for reserve actuator moments, particularly in distal limb joints. The high sensitivity in distal limb muscles to TSL changes indicated that reserve actuators were directly compensating for the reduced muscle capacity to generate the moments required for STS and STW. Simulations incorporating full tissue properties notably reduced the necessity for reserve actuators at distal limb joints in both movements, highlighting the crucial role of passive tissues during standing up. It was notable that although STW required larger ankle flexion/extension joint moments than STS, the percentages of ankle reserve actuator moments compared to net joint moments were smaller, suggesting a more effective use of muscles contributing to ankle extension. However, STW required larger percentages of reserve actuator moments at the hip and knee joints.

### Sensitivity analyses

Static optimisation simulation results were influenced by ± 5 % TSL changes for numerous muscles, particularly when the ratio of muscle fibre length to tendon length was smaller (**Figures 6 – 9**, dashed and dotted lines). Hip and knee muscles showed lower sensitivity to TSL changes, although activations and forces early in the transition were impacted. Patterns for normalised fibre lengths for specific muscles like PIFML and FMTI were moderately shifted closer to their optimal length limit (for shorter tendons) or moved toward/below the optimal length (for longer tendons). In contrast, ankle and digit muscles displayed substantial changes in activation, length patterns, and forces. For instance, with a decreased TSL, FL reached its maximum possible activation (1.0) throughout both transitions, while FL, FP3, and FPP3 dropped to zero activation when TSL was increased. GL and GM moderately increased activations with an increased TSL, reflecting complex force patterns. GM notably generated higher force (∼ 6 x BW) early in both transitions when TSL was increased, while FP3 and FPP3 showed minimal force across all TSL adjustments. These changes were reflected in the normalised fibre lengths in the same ways as for proximal muscles, where decreased TSL values led to longer fibre lengths.

In both STS and STW, additional dynamic simulations revealed that simulations with compliant tendons but no passive muscle tissue yielded results akin to simulations with full tissue properties, while dynamic simulations with all tissue properties limited showed similar muscle activation patterns and forces as static simulations, but slightly reduced reserve actuator requirements (**Table S2**). This result is consistent with the inference that tendons play a substantial role in muscle activation, force generation, and fibre operating lengths compared to other passive tissues.

Static optimisation simulations of the second set of trials (one each for STS and STW) had some changes in predicted limb muscle activations (**Figure S4**). In the second STS trial, activation levels were generally lower for muscles (including CFP which showed minimal activations) except for a few muscles such as GI and FPP3. Several muscles showed two peak activations during the transition such as ILPOp, which was activated in both the flexion momentum phase (before heel-off) and the ascending phase (after heel-off). In the second STW trial, the activation levels were similar, but some hip and knee muscles showed high activations late in the ascending phase, such as ILPR, in comparison with the first trial where those muscles were mostly activated early in the transition (around heel-off). The unusual GM activity and force in the original simulation were not present when using this alternative trial. Overall, while some quantitative results varied between the trials, qualitative patterns were broadly similar, such as high activations in major extensor muscles; which peaked around heel-off; and extreme fibre length changes, especially for distal muscles.

## Discussion

Our study provides the first dataset of hindlimb kinematics and kinetics as well as simulated muscle dynamics during emu sit-to-stand and sit-to-walk transitions; or for any bird. We posited four hypotheses concerning emu hindlimb muscle activations, fibre length changes, passive tissue roles, and differences in muscular demands between STS and STW. Overall, our findings support all four hypotheses.

Supporting Hypothesis 1, our findings revealed substantial activations and forces in many extensor muscles as well as key muscles contributing to non-sagittal movements during both STS and STW (**Figures 6 – 9**). Major hip, knee, and ankle extensors such as PIFML, ILPR, FMTI, GL, and GM had the greatest activations and forces. Although emus primarily required flexor moments at the knee joints during the movements, the knee extensor muscles were likely activated to counter the knee flexion moments generated by ankle extensors (e.g., GL and FPP3). Moreover, we found that muscles transitioning from initially sub-optimal fibre lengths (e.g., GM) were preferred for force generation over those operating closer to optimal length values (e.g., OMII) as demonstrated by their higher force generation. Co-activations of medial (e.g., ITCr and ITCaa) and lateral rotators (e.g., OMII and OMIP) were also observed, essential for opposing the adduction of the hindlimb by the GRF as a result of the quasi-parasagittal gait in birds (Gatesy, 1994, 1999; Hutchinson and Gatesy, 2000).

Hypothesis 2 postulated that emu hindlimb muscles would operate near their functional limits. Unlike the more isometric muscle patterns observed in forward locomotion (e.g., Bishop *et al*., 2021), our nominal static simulation results showed extensive fibre length changes during standing up, closely approaching their functional limits (0.5 – 1.5 times optimal fibre length), especially in distal muscles (**Figures 6 – 9**). Some muscles such as GL and FP3 were further away from this limit but exhibited higher sensitivity to changes in TSL and simulation frameworks. In our model building approach, we systematically adjusted TSLs and optimal fibre in order to keep muscle fibres within a reasonable operating range. While this process could have biased the analysis towards (or even away from) extreme muscle lengths, our modifications were cautious, ensuring muscles avoided excessive lengthening or shortening. Thus, we contend that the approach was appropriate.

Hypothesis 3 proposed the necessity of passive support mechanisms (namely tendons), especially in the distal hindlimbs, for executing STS and STW tasks. Modifying TSL by ± 5% notably amplified the demand for reserve actuator moments, especially in the distal hindlimbs, where muscle fibres approached their functional limits (**Table S2**). This outcome was not surprising, considering that muscles operating beyond their optimal length ranges encounter difficulties in generating adequate forces (e.g., Zajac, 1989; Millard *et al*., 2013). Muscle fibre length changes reduced in simulations incorporating full tissue properties compared to nominal static simulations, especially for distal muscles (**Figures 6 – 9**). There was also a general reduction in muscle forces, with the exception of a few ankle extensors (e.g., GM and FL) (**Figures 6 – 9**). This discrepancy may be attributed to the increased force generation capacity of the muscles allowing for improved coordination (reduced co-contraction) of other muscles acting around the same DOF (e.g., GL, FP3, and FPP3), while the total required joint moments remained constant. Subsequent investigations that quantify muscle work could contribute to a clearer understanding of these dynamics. Overall, our results indicated the crucial requirement for passive support around various DOFs, particularly for distal hindlimb joints. This aligns with walking and running simulations of other birds, which have also highlighted the necessity of passive support in non-sagittal motions and distal limb joints (Rankin *et al*., 2016; Bishop, Michel, *et al*., 2021).

Hypothesis 4 proposed that the STW transition would involve greater demands on muscle coordination than the STS transition in emus, predominantly for extensor muscles and non-sagittal muscles. We found that STW involved greater non-sagittal motions at the hip and knee than in STS (**Table 1**, **Figure 3**). Consequently, in emus, STW muscle coordination may be more demanding than STS in order to meet the demands of non-sagittal joint moments. Simulation results support this notion, demonstrating that STW generally required higher activations and muscle forces than STS (**Figures 6 – 9**). STW also required reserve actuator moments that were greater for hip and knee joints compared to STS, but lower for the ankle plantarflexion/dorsiflexion DOF (**Table S2**). Our results suggest compensatory strategies may be used in STW to spare the load of ankle extensors with elevated hip and knee muscle activities, discussed below.

The movement patterns of emu STS and STW behaviours exhibited similarities and distinctions when compared to humans. Emus demonstrated a sequential joint excursion from proximal to distal during the transition like humans do (Pandy, Garner and Anderson, 1995), an observation also seen in greyhounds (Ellis, Rankin and Hutchinson, 2018). Both emu movements involved generating substantial forward momentum, positioning the COM behind the toes at heel-off, resembling the ‘momentum transfer’ or ‘forward momentum’ strategy observed in humans (Hughes, 1996; Norman-Gerum and McPhee, 2020; Perera *et al*., 2023). However, it is important to note the differences between emus and humans in performing the two tasks: while human STW demands greater stability, often with a longer flexion (or pitch) momentum period sacrificing speed or efficiency (Kerr, Durward and Kerr, 2004), emus had a relatively small body pitch movement and displayed a faster flexion momentum period in STW than in STS (**Table 1**). Emus also seem to have a relatively smaller braking impulse compared to humans during the transition (e.g., Magnan, McFadyen and St-Vincent, 1996; Perera et al., 2023), especially when performing the STW task (Table 1). The braking impulse reduces forward momentum and presumably allows a focus on stability and postural control when standing up (Perera et al., 2023). These differences suggest that emus transition from sitting to walking while maintaining stability, possibly due to their cranially positioned COM and use of passive support mechanisms. This speculation remains untested but aligns with hypotheses on birds’ energy-efficient stabilisation methods (Abourachid *et al*., 2023). Further support for this idea comes from the smaller medio-lateral COP (Table 1) – regarded as a balance indicator (Thompson et al., 2017) – observed in STW relative to STS.

While direct measurements of emu hindlimb muscle forces and electromyographic (EMG) data were unavailable, simulation results suggested that emus had many muscle recruitment patterns similar to humans (Pandy *et al*., 1995), with a few noticeable differences. In the dynamic simulations (**Figures 7 – 10**), muscles generally followed a proximal-to-distal activation sequence, gradually recruiting proximal muscles (e.g., PIFML, ILPOp) until a more upright posture was achieved, with distal muscles (e.g., FP3, FPP3) activated later. However, a few muscles (e.g., GM) remained continuously active. Static optimisation simulations did not exhibit this obvious activation sequence, likely due to limitations in our static optimisation framework, as observed in previous research (Ellis *et al*., 2018). Across all simulations, we observed co-contractions of antagonist muscles, likely crucial for redistribution of joint moments and stability (e.g., Roebroeck *et al*., 1994; Khemlani, Carr and Crosbie, 1999; Savelberg *et al*., 2007; Actis *et al*., 2018; Shia *et al*., 2018). In contrast to humans, emus required a relatively small net extensor moment at the knee joint (**Figure 5**), likely a result of their more horizontally oriented trunk and anteriorly positioned COM. This adaptation, however, would potentially increase the load on ankle extensors. This redistribution of moments becomes particularly evident in STW, characterised by increased net joint moments at the hip and ankle joints, with increased trunk flexion and a more forward position of the point of application of the GRF (**Figure 5**). The trade-offs between fibre lengths and capacity pose additional challenges for ankle extensors, suggesting that the ankle extensor capacity represents a key biomechanical constraint in cursorial species during standing up, as well as potentially in running (Hutchinson, 2004).

How do emus navigate these biomechanical challenges during standing up? Our study suggests that emus rely on some key pelvic limb muscles other than extensors, use substantial movement in non-sagittal planes, and leverage the elastic properties of their tendons to forestall large fibre length changes as well as muscle forces, particularly in distal muscles – an observation consistent with previous research on greyhounds (Ellis *et al*., 2018). However, the efficacy of tendons in elastic energy storage in STS/STW might be limited compared to their roles in running (e.g., Alexander *et al*., 1979; Roberts *et al*., 1998, p. 199; Rubenson *et al*., 2011) and jumping (e.g., Henry, Ellerby and Marsh, 2005; Konow and Roberts, 2015; Bishop *et al*., 2021). The challenge here is that muscles need to maintain quasi-isometric conditions to effectively recover strain energy when tendons recoil (Biewener, 1998). Elastic energy storage depends on rapid loading and unloading, differing from the demands of standing up.

This study rested on two pivotal assumptions: bilateral symmetry and the roles of tendons. Although we assumed bilateral symmetry during the ‘flexion momentum’ and ‘ascending’ phases of the STS and STW transitions, asymmetrical movements or forces between contralateral limbs across different planes were likely, an observation also seen in human STS and STW studies (e.g., Lundin, Grabiner and Jahnigen, 1995; Boukadida *et al*., 2015; Dolecka, Ownsworth and Kuys, 2015; Caruthers *et al*., 2016). This asymmetry might serve as a compensatory mechanism during forward acceleration of the COM (van der Kruk *et al*., 2021) or for quicker balance recovery (Blaszczyk *et al*., 2000). In addition, we expected that dynamic simulations would highlight tendons’ roles in preventing large muscle fibre forces, activations, and length changes. However, simplifications in our model might limit a comprehensive understanding. For example, our model did not include other passive tissues, such as ligaments, which are considered to be highly specialised for energy savings in large ratites (e.g., Schaller, 2008; Schaller *et al*., 2009; Badri-Spröwitz *et al*., 2022). The stretch-shortening effect of muscles (Anderson and Pandy, 1993; Daley and Biewener, 2003; Bobbert and Casius, 2005) could also play a role during standing up in some muscles such as GL, which underwent active lengthening and then shortening. Tuning TSLs to maintain muscle fibre lengths within a feasible range presents another relevant challenge, which currently lacks a standardised approach (e.g., Scovil and Ronsky, 2006; Redl, Gfoehler and Pandy, 2007; Bishop *et al*., 2021).

The calculation of joint angles using skin markers and the estimation of single limb GRFs from combined GRF data in our study come with inherent limitations that should also be acknowledged. Joint kinematics computed from skin markers can be substantially influenced by factors such as calibration accuracy (Chiari *et al*., 2005), skin movement (Leardini *et al*., 2005) and marker placement errors (Della Croce *et al*., 2005). The criteria for measurement accuracy also was derived originally from human studies (Hicks *et al*., 2015). Despite our best efforts to account for these challenges, we still observed large variations in knee and ankle rotation angles, potentially attributed to errors from the placement of tibiotarsus markers (**Figures S1**). The partitioning of GRFs based on the assumption of bilateral symmetry may lead to an underestimation of actual GRFs, particularly notable for the stance limb during the STW transition. Additionally, imprecise COP positions could also influence inverse dynamics moments. While a more comprehensive procedure would improve accuracy (e.g., Rubenson *et al*., 2007), it would require new data collection and reanalysis of newly generated simulations. Alternative methods, such as tracking simulations that partly account for error in experimental data, could mitigate inconsistencies between model kinematics and kinetics (e.g., Bishop *et al*., 2021). Incorporating upper body movement (including head and neck) into our model may yield additional insights into STS and STW, as observed in humans who use arm movements for standing up (Mazzà *et al*., 2004; Davidson *et al*., 2013; Dolecka *et al*., 2015; Komaris *et al*., no date). Although emus seem to minimally use such movements during standing up, their apparently extensive neck movements during STS might contribute to generating forward momentum and shifting the COM closer to the BOS. Emus seemed to use fewer neck movements during STW, perhaps relying more on trunk movements to generate horizontal momentum (**Videos S1 and S2**), suggesting distinct constraints and strategies between humans and emus in standing up.

## Conclusions

As the first investigation on sit-to-stand and sit-to-walk transitions in an avian species (or even among very few for non-humans in general), this study unravels joint mechanics, constraints, and compensatory strategies used by emus during these movements. Emus demonstrate large muscle activations, substantial muscle force requirements, and fibre length changes, particularly in their distal hindlimbs, with ankle extensors acting as a key biomechanical limit during standing up. To mitigate trade-offs between muscle capacity and fibre lengths, emus use non-sagittal muscle actions and tendon length changes, which is more evident in transitioning from sitting to walking. The current study lays a groundwork for broader investigations spanning diverse species. Understanding the foundational biomechanics of sit-to-stand and sit-to-walk transitions in animals holds promise for insights into morphofunctional specialisations, body size influences, evolutionary studies, applications in robotics, and advancements in animal welfare.

## Acknowledgements

We thank Andrea S. Pollard and staff of the Structure and Motion Laboratory for assistance with data collection. We thank James Usherwood, Monica A. Daley, and Pasha van Bijlerts for helpful advice and discussions. YL is very appreciative of Delyle T. Polet, Masaya Iijima, Mauro B. C. Lacerda, and Stacy Ashlyn for sharing office space and engaging in discussions on mechanics, data processing, and musculoskeletal modelling and simulations in general.

## Data availability

The musculoskeletal model and motion data used in this study have been provided in Supplementary Data S1. Supplementary texts and figures are provided as a separate document.

We have also produced three supplementary videos:

Video S1 – The exemplar trial of emu sit-to-stand transition.

Video S2 – The exemplar trial of emu sit-to-walk transition.

Video S3 – Animation of emu sit-to-stand and sit-to-walk transitions using inverse dynamics simulations with full tissue properties. Video played at 0.2 x original speed.

## List of symbols and abbreviations

*a*: Simulated muscle activation
*F_max_*: Muscle’s maximal isometric force
*F_m_\**: Body weight-nondimensionalised muscle force
*l\**: Normalised fibre length
*m*: Muscle belly mass
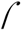: Muscle’s maximum isometric stress
*〉*: Muscle tissue density
3D: Three-dimensional
AMB: M. ambiens
BOS: Base of support
BW: Body weight
CFP: M. caudofemoralis p. pelvica
COM: Centre of mass
COP: Centre of pressure
CT: Computed tomography
DOF: Degrees of freedom
EDL: M. extensor digitorum longus
EMA: Effective mechanical advantage
EMG: Electromyography
FCLA: M. flexor cruris lateralis pars accessoria
FCLP: M. flexor cruris lateralis pars pelvica
FCM: M. flexor cruris medialis
FDL: M. flexor digitorum longus
FHL: M. flexor hallucis longus
FL: M. fibularis longus
FMTI: M. femorotibialis intermedialis
FMTL: M. femorotibialis lateralis
FMTM: M. femorotibialis medialis
FP2: M. flexor perforatus digiti II
FP3: M. flexor perforatus digiti III
FP4: M. flexor perforatus digiti IV
FPP2: M. flexor perforans et perforatus digiti II
FPP3: M. flexor perforans et perforatus digiti III
GI: M. gastrocnemius (pars) intermedius
GL: M. gastrocnemius (pars) lateralis
GM: M. gastrocnemius (pars) medialis
GRF: Ground reaction force
IC: M. iliotibialis cranialis
IFE: M. iliofemoralis externus
IFI: M. iliofemoralis internus
ILFB: M. iliofibularis
ILPOa: M. iliotibialis lateralis pars postacetabularis (anterior part)
ILPOp: M. iliotibialis lateralis pars postacetabularis (posterior part)
ILPR: M. iliotibialis lateralis pars preacetabularis
ISF: M. ischiofemoralis
ITCaa: M. iliotrochantericus caudalis (anterior part)
ITCap: M. iliotrochantericus caudalis (posterior part)
ITCr: M. iliotrochantericus cranialis
ITM: M. iliotrochantericus medialis
ML: Medio-lateral
MTP: Tarsometatarsophalangeal
MTU: Muscle-tendon unit
OMII: M. obturatorius medialis (Ilium-ischium part)
OMIP: M. obturatorius medialis (Ischium-pubis part)
PIFML: M. puboischiofemoralis p. lateralis and p. medialis
ROM: Range of motion
SD: Standard deviation
STS: Sit-to-stand
STW: Sit-to-walk
TCf: M. tibialis cranialis caput femorale
TCt: M. tibialis cranialis caput tibiale
TSL: Tendon slack length

## References

Abourachid, A., Chevallereau, C., Pelletan, I. and Wenger, P. (2023) ‘An upright life, the postural stability of birds: a tensegrity system’, Journal of The Royal Society Interface, 20(208), p. 20230433. Available at: 10.1098/rsif.2023.0433.

Actis, J.A., Nolasco, L.A., Gates, D.H. and Silverman, A.K. (2018) ‘Lumbar loads and trunk kinematics in people with a transtibial amputation during sit-to-stand’, Journal of Biomechanics, 69, pp. 1–9. Available at: 10.1016/j.jbiomech.2017.12.030.

Aissaoui, R. and Dansereau, J. (1999) ‘Biomechanical analysis and modelling of sit to stand task: a literature review’, in IEEE SMC’99 Conference Proceedings. 1999 IEEE International Conference on Systems, Man, and Cybernetics (Cat. No.99CH37028). IEEE SMC’99 Conference Proceedings. 1999 IEEE International Conference on Systems, Man, and Cybernetics (Cat. No.99CH37028), pp. 141–146 vol.1. Available at: 10.1109/ICSMC.1999.814072.

Alexander, R.McN., Maloiy, G.M.O., Njau, R. and Jayes, A.S. (1979) ‘Mechanics of running of the ostrich (Struthio camelus)’, Journal of Zoology, 187(2), pp. 169–178. Available at: 10.1111/j.1469-7998.1979.tb03941.x.

Anderson, F.C. and Pandy, M.G. (1993) ‘Storage and utilization of elastic strain energy during jumping’, Journal of Biomechanics, 26(12), pp. 1413–1427. Available at: 10.1016/0021-9290(93)90092-S.

Arnold, E.M. and Delp, S.L. (2011) ‘Fibre operating lengths of human lower limb muscles during walking’, Philosophical Transactions of the Royal Society B: Biological Sciences, 366(1570), pp. 1530–1539. Available at: 10.1098/rstb.2010.0345.

Badri-Spröwitz, A., Aghamaleki Sarvestani, A., Sitti, M. and Daley, M.A. (2022) ‘BirdBot achieves energy-efficient gait with minimal control using avian-inspired leg clutching’, Science Robotics, 7(64), p. eabg4055. Available at: 10.1126/scirobotics.abg4055.

Biewener, A.A. (1989) ‘Scaling Body Support in Mammals: Limb Posture and Muscle Mechanics’, Science, 245(4913), pp. 45–48. Available at: 10.1126/science.2740914.

Biewener, A.A. (1998) ‘Muscle Function in vivo: A Comparison of Muscles Used for Elastic Energy Savings versus Muscles Used to Generate Mechanical Power’, American Zoologist, 38(4), pp. 703–717. Available at: doi10.1093/icb/38.4.703.

Biewener, A.A. (2005) ‘Biomechanical consequences of scaling’, Journal of Experimental Biology, 208(9), pp. 1665–1676. Available at: 10.1242/jeb.01520.

Bishop, P.J., Falisse, A., De Groote, F. and Hutchinson, J.R. (2021) ‘Predictive Simulations of Musculoskeletal Function and Jumping Performance in a Generalized Bird’, Integrative Organismal Biology, 3(1), p. obab006. Available at: 10.1093/iob/obab006.

Bishop, P.J., Michel, K.B., Falisse, A., Cuff, A.R., Allen, V.R., Groote, F.D. and Hutchinson, J.R. (2021) ‘Computational modelling of muscle fibre operating ranges in the hindlimb of a small ground bird (Eudromia elegans), with implications for modelling locomotion in extinct species’, PLOS Computational Biology, 17(4), p. e1008843. Available at: 10.1371/journal.pcbi.1008843.

Bishop, P.J., Wright, M.A. and Pierce, S.E. (2021) ‘Whole-limb scaling of muscle mass and force-generating capacity in amniotes’, PeerJ, 9, p. e12574. Available at: 10.7717/peerj.12574.

Blaszczyk, J.W., Prince, F., Raiche, M. and Hébert, R. (2000) ‘Effect of ageing and vision on limb load asymmetry during quiet stance’, Journal of Biomechanics, 33(10), pp. 1243–1248. Available at: 10.1016/S0021-9290(00)00097-X.

Bobbert, M.F. and Casius, L.J.R. (2005) ‘Is the Effect of a Countermovement on Jump Height due to Active State Development?’, Medicine & Science in Sports & Exercise, 37(3), p. 440. Available at: 10.1249/01.MSS.0000155389.34538.97.

Bobbert, M.F., Kistemaker, D.A., Vaz, M.A. and Ackermann, M. (2016) ‘Searching for strategies to reduce the mechanical demands of the sit-to-stand task with a muscle-actuated optimal control model’, Clinical Biomechanics, 37, pp. 83–90. Available at: 10.1016/j.clinbiomech.2016.06.008.

Boukadida, A., Piotte, F., Dehail, P. and Nadeau, S. (2015) ‘Determinants of sit-to-stand tasks in individuals with hemiparesis post stroke: A review’, Annals of Physical and Rehabilitation Medicine, 58(3), pp. 167–172. Available at: 10.1016/j.rehab.2015.04.007.

Brouwers, S.P., Simmler, M., Savary, P. and Scriba, M.F. (2023) ‘Towards a novel method for detecting atypical lying down and standing up behaviors in dairy cows using accelerometers and machine learning’, Smart Agricultural Technology, 4, p. 100199. Available at: 10.1016/j.atech.2023.100199.

Carrano, M.T. (1999) ‘What, if anything, is a cursor? Categories versus continua for determining locomotor habit in mammals and dinosaurs’, Journal of Zoology, 247(1), pp. 29–42. Available at: 10.1111/j.1469-7998.1999.tb00190.x.

Caruthers, E.J., Thompson, J.A., Chaudhari, A.M.W., Schmitt, L.C., Best, T.M., Saul, K.R. and Siston, R.A. (2016) ‘Muscle Forces and Their Contributions to Vertical and Horizontal Acceleration of the Center of Mass During Sit-to-Stand Transfer in Young, Healthy Adults’, Journal of Applied Biomechanics, 32(5), pp. 487–503. Available at: 10.1123/jab.2015-0291.

Chiari, L., Croce, U.D., Leardini, A. and Cappozzo, A. (2005) ‘Human movement analysis using stereophotogrammetry: Part 2: Instrumental errors’, Gait & Posture, 21(2), pp. 197–211. Available at: 10.1016/j.gaitpost.2004.04.004.

Daley, M.A. and Biewener, A.A. (2003) ‘Muscle force-length dynamics during level versus incline locomotion: a comparison of in vivo performance of two guinea fowl ankle extensors’, Journal of Experimental Biology, 206(17), pp. 2941–2958. Available at: 10.1242/jeb.00503.

Davidson, B.S., Judd, D.L., Thomas, A.C., Mizner, R.L., Eckhoff, D.G. and Stevens-Lapsley, J.E. (2013) ‘Muscle activation and coactivation during five-time-sit-to-stand movement in patients undergoing total knee arthroplasty’, Journal of Electromyography and Kinesiology, 23(6), pp. 1485–1493. Available at: 10.1016/j.jelekin.2013.06.008.

De Groote, F., Kinney, A.L., Rao, A.V. and Fregly, B.J. (2016) ‘Evaluation of Direct Collocation Optimal Control Problem Formulations for Solving the Muscle Redundancy Problem’, Annals of Biomedical Engineering, 44(10), pp. 2922–2936. Available at: 10.1007/s10439-016-1591-9.

Dehail, P., Bestaven, E., Muller, F., Mallet, A., Robert, B., Bourdel-Marchasson, I. and Petit, J. (2007) ‘Kinematic and electromyographic analysis of rising from a chair during a “Sit-to-Walk” task in elderly subjects: Role of strength’, Clinical Biomechanics, 22(10), pp. 1096– 1103. Available at: 10.1016/j.clinbiomech.2007.07.015.

Della Croce, U., Leardini, A., Chiari, L. and Cappozzo, A. (2005) ‘Human movement analysis using stereophotogrammetry: Part 4: assessment of anatomical landmark misplacement and its effects on joint kinematics’, Gait & Posture, 21(2), pp. 226–237. Available at: 10.1016/j.gaitpost.2004.05.003.

Delp, S.L., Anderson, F.C., Arnold, A.S., Loan, P., Habib, A., John, C.T., Guendelman, E. and Thelen, D.G. (2007) ‘OpenSim: Open-Source Software to Create and Analyze Dynamic Simulations of Movement’, IEEE Transactions on Biomedical Engineering, 54(11), pp. 1940–1950. Available at: 10.1109/TBME.2007.901024.

Dembia, C.L., Bianco, N.A., Falisse, A., Hicks, J.L. and Delp, S.L. (2020) ‘OpenSim Moco: Musculoskeletal optimal control’, PLOS Computational Biology, 16(12), p. e1008493. Available at: 10.1371/journal.pcbi.1008493.

Dick, T.J.M. and Clemente, C.J. (2017) ‘Where Have All the Giants Gone? How Animals Deal with the Problem of Size’, PLOS Biology, 15(1), p. e2000473. Available at: 10.1371/journal.pbio.2000473.

Dolecka, U.E., Ownsworth, T. and Kuys, S.S. (2015) ‘Comparison of sit-to-stand strategies used by older adults and people living with dementia’, Archives of Gerontology and Geriatrics, 60(3), pp. 528–534. Available at: 10.1016/j.archger.2014.12.007.

Doorenbosch, C.A.M., Harlaar, J., Roebroeck, M.E. and Lankhorst, G.J. (1994) ‘Two strategies of transferring from sit-to-stand; The activation of monoarticular and biarticular muscles’, Journal of Biomechanics, 27(11), pp. 1299–1307. Available at: 10.1016/0021-9290(94)90039-6.

Ellis, M.I., Seedhom, B.B. and Wright, V. (1984) ‘Forces in the knee joint whilst rising from a seated position’, Journal of Biomedical Engineering, 6(2), pp. 113–120. Available at: 10.1016/0141-5425(84)90053-0.

Ellis, R.G., Rankin, J.W. and Hutchinson, J.R. (2018) ‘Limb Kinematics, Kinetics and Muscle Dynamics During the Sit-to-Stand Transition in Greyhounds’, Frontiers in Bioengineering and Biotechnology, 6. Available at: https://www.frontiersin.org/articles/10.3389/fbioe.2018.00162 (Accessed: 17 December 2022).

Feeney, L.C., Lin, C.-F., Marcellin-Little, D.J., Tate, A.R., Queen, R.M. and Yu, B. (2007) ‘Validation of two-dimensional kinematic analysis of walk and sit-to-stand motions in dogs’, American Journal of Veterinary Research, 68(3), pp. 277–282. Available at: 10.2460/ajvr.68.3.277.

Fukunaga, T., Kubo, K., Kawakami, Y., Fukashiro, S., Kanehisa, H. and Maganaris, C.N. (2001) ‘In vivo behaviour of human muscle tendon during walking’, Proceedings of the Royal Society of London. Series B: Biological Sciences, 268(1464), pp. 229–233. Available at: 10.1098/rspb.2000.1361.

Gardner, R.B. (2011) ‘Evaluation and Management of the Recumbent Adult Horse’, Veterinary Clinics of North America: Equine Practice, 27(3), pp. 527–543. Available at: 10.1016/j.cveq.2011.08.006.

Gatesy, S.M. (1994) ‘Neuromuscular diversity in archosaur deep dorsal thigh muscles’, Brain, Behavior and Evolution, 43(1), pp. 1–14. Available at: 10.1159/000113619.

Gatesy, S.M. (1999) ‘Guineafowl hind limb function. I: Cineradiographic analysis and speed effects’, Journal of Morphology, 240(2), pp. 115–125. Available at: 10.1002/(SICI)1097-4687(199905)240:2<115::AID-JMOR3>3.0.CO;2-Y.

Goetz, J.E., Derrick, T.R., Pedersen, D.R., Robinson, D.A., Conzemius, M.G., Baer, T.E. and Brown, T.D. (2008) ‘Hip joint contact force in the emu (Dromaius novaehollandiae) during normal level walking’, Journal of Biomechanics, 41(4), pp. 770–778. Available at: 10.1016/j.jbiomech.2007.11.022.

Henry, H.T., Ellerby, D.J. and Marsh, R.L. (2005) ‘Performance of guinea fowl Numida meleagris during jumping requires storage and release of elastic energy’, Journal of Experimental Biology, 208(17), pp. 3293–3302. Available at: 10.1242/jeb.01764.

Hicks, J.L., Uchida, T.K., Seth, A., Rajagopal, A. and Delp, S.L. (2015) ‘Is My Model Good Enough? Best Practices for Verification and Validation of Musculoskeletal Models and Simulations of Movement’, Journal of Biomechanical Engineering, 137(2). Available at: 10.1115/1.4029304.

Hortobágyi, T., Mizelle, C., Beam, S. and DeVita, P. (2003) ‘Old Adults Perform Activities of Daily Living Near Their Maximal Capabilities’, The Journals of Gerontology: Series A, 58(5), pp. M453–M460. Available at: 10.1093/gerona/58.5.M453.

Hughes, M.A. (1996) ‘Chair rise strategy in the functionally impaired elderly’. Available at: doi10.1016/S0021-9290(96)80001-7.

Hughes, M.A., Weiner, D.K., Schenkman, M.L., Long, R.M. and Studenski, S.A. (1994) ‘Chair rise strategies in the elderly’, Clinical Biomechanics, 9(3), pp. 187–192. Available at: 10.1016/0268-0033(94)90020-5.

Hutchinson, J.R. (2004) ‘Biomechanical modeling and sensitivity analysis of bipedal running ability. I. Extant taxa’, Journal of Morphology, 262(1), pp. 421–440. Available at: 10.1002/jmor.10241.

Hutchinson, J.R. and Gatesy, S.M. (2000) ‘Adductors, Abductors, and the Evolution of Archosaur Locomotion’, Paleobiology, 26(4), pp. 734–751. Available at: doi:10.1666/0094-8373(2000)026<0734:AAATEO>2.0.CO;2.

Hutchinson, J.R., Rankin, J.W., Rubenson, J., Rosenbluth, K.H., Siston, R.A. and Delp, S.L. (2015) ‘Musculoskeletal modelling of an ostrich (Struthio camelus) pelvic limb: influence of limb orientation on muscular capacity during locomotion’, PeerJ, 3, p. e1001. Available at: 10.7717/peerj.1001.

Kerr, A., Durward, B. and Kerr, K.M. (2004) ‘Defining phases for the sit-to-walk movement’, Clinical Biomechanics, 19(4), pp. 385–390. Available at: 10.1016/j.clinbiomech.2003.12.012.

Khemlani, M.M., Carr, J.H. and Crosbie, W.J. (1999) ‘Muscle synergies and joint linkages in sit-to-stand under two initial foot positions’, Clinical Biomechanics, 14(4), pp. 236–246. Available at: 10.1016/S0268-0033(98)00072-2.

Komaris, D.-S., Govind, C., Murphy, A., Ewen, A. and Riches, P. (no date) ‘Identification of Movement Strategies During the Sit-to-Walk Movement in Patients With Knee Osteoarthritis’, Journal of Applied Biomechanics, 34(2), pp. 96–103. Available at: 10.1123/jab.2016-0279.

Konow, N. and Roberts, T.J. (2015) ‘The series elastic shock absorber: tendon elasticity modulates energy dissipation by muscle during burst deceleration’, Proceedings of the Royal Society B: Biological Sciences, 282(1804), p. 20142800. Available at: 10.1098/rspb.2014.2800.

Kralj, A., Jaeger, R.J. and Munih, M. (1990) ‘Analysis of standing up and sitting down in humans: Definitions and normative data presentation’, Journal of Biomechanics, 23(11), pp. 1123–1138. Available at: 10.1016/0021-9290(90)90005-N.

van der Kruk, E., Silverman, A.K., Reilly, P. and Bull, A.M.J. (2021) ‘Compensation due to age-related decline in sit-to-stand and sit-to-walk’, Journal of Biomechanics, 122, p. 110411. Available at: 10.1016/j.jbiomech.2021.110411.

Lamas, L.P., Main, R.P. and Hutchinson, J.R. (2014) ‘Ontogenetic scaling patterns and functional anatomy of the pelvic limb musculature in emus (Dromaius novaehollandiae)’, PeerJ, 2, p. e716. Available at: 10.7717/peerj.716.

Lamas, L.R.G.P. (2015) Musculoskeletal biomechanics during growth on emu (Dromaius; Aves) : an integrative experimental and modelling analysis. Ph.D. University of London Royal Veterinary College. Available at: https://ethos.bl.uk/OrderDetails.do?uin=uk.bl.ethos.701660 (Accessed: 13 June 2023).

Leardini, A., Chiari, L., Croce, U.D. and Cappozzo, A. (2005) ‘Human movement analysis using stereophotogrammetry: Part 3. Soft tissue artifact assessment and compensation’, Gait & Posture, 21(2), pp. 212–225. Available at: 10.1016/j.gaitpost.2004.05.002.

Lichtwark, G.A., Bougoulias, K. and Wilson, A.M. (2007) ‘Muscle fascicle and series elastic element length changes along the length of the human gastrocnemius during walking and running’, Journal of Biomechanics, 40(1), pp. 157–164. Available at: 10.1016/j.jbiomech.2005.10.035.

Lichtwark, G.A. and Wilson, A.M. (2006) ‘Interactions between the human gastrocnemius muscle and the Achilles tendon during incline, level and decline locomotion’, Journal of Experimental Biology, 209(21), pp. 4379–4388. Available at: 10.1242/jeb.02434.

Lidfors, L. (1989) ‘The use of getting up and lying down movements in the evaluation of cattle environments’, Veterinary Research Communications, 13(4), pp. 307–324. Available at: 10.1007/BF00420838.

Lundin, T.M., Grabiner, M.D. and Jahnigen, D.W. (1995) ‘On the assumption of bilateral lower extremity joint moment symmetry during the sit-to-stand task’, Journal of Biomechanics, 28(1), pp. 109–112. Available at: 10.1016/0021-9290(95)80013-1.

Maloiy, G.M.O., Alexander, R.McN., Njau, R. and Jayes, A.S. (1979) ‘Allometry of the legs of running birds’, Journal of Zoology, 187(2), pp. 161–167. Available at: 10.1111/j.1469-7998.1979.tb03940.x.

Manal, K. and Buchanan, T. (2004) ‘Subject-Specific Estimates of Tendon Slack Length: A Numerical Method’, Journal of Applied Biomechanics, 20, pp. 195–203. Available at: 10.1123/jab.20.2.195.

Mazzà, C., Benvenuti, F., Bimbi, C. and Stanhope, S.J. (2004) ‘Association between subject functional status, seat height, and movement strategy in sit-to-stand performance’, Journal of the American Geriatrics Society, 52(10), pp. 1750–1754. Available at: 10.1111/j.1532-5415.2004.52472.x.

Medler, S. (2002) ‘Comparative trends in shortening velocity and force production in skeletal muscles’, *American Journal of Physiology-Regulatory*, Integrative and Comparative Physiology, 283(2), pp. R368–R378. Available at: 10.1152/ajpregu.00689.2001.

Mendez, J. and Keys, A. (1960) ‘Density and composition of mammalian muscle’, Metabolism, Clinical and Experimental, 9(2), pp. 184–188.

Millard, M., Uchida, T., Seth, A. and Delp, S.L. (2013) ‘Flexing Computational Muscle: Modeling and Simulation of Musculotendon Dynamics’, Journal of Biomechanical Engineering, 135(2). Available at: 10.1115/1.4023390.

Nina U. Schaller (2008) Structural attributes contributing to locomotor performance in the ostrich. University of Heidelberg. Available at: https://archiv.ub.uni-heidelberg.de/volltextserver/8852/1/Schaller_Dis.pdf (Accessed: 7 December 2023).

Norman-Gerum, V. and McPhee, J. (2020) ‘Comprehensive description of sit-to-stand motions using force and angle data’, Journal of Biomechanics, 112, p. 110046. Available at: 10.1016/j.jbiomech.2020.110046.

Pandy, M.G., Garner, B.A. and Anderson, F.C. (1995) ‘Optimal Control of Non-ballistic Muscular Movements: A Constraint-Based Performance Criterion for Rising From a Chair’, Journal of Biomechanical Engineering, 117(1), pp. 15–26. Available at: 10.1115/1.2792265.

Perera, C.K., Gopalai, A.A., Gouwanda, D., Ahmad, S.A. and Salim, M.S.B. (2023) ‘Sit-to-walk strategy classification in healthy adults using hip and knee joint angles at gait initiation’, Scientific Reports, 13, p. 16640. Available at: 10.1038/s41598-023-43148-0.

Rankin, J.W., Rubenson, J. and Hutchinson, J.R. (2016) ‘Inferring muscle functional roles of the ostrich pelvic limb during walking and running using computer optimization’, Journal of The Royal Society Interface, 13(118), p. 20160035. Available at: 10.1098/rsif.2016.0035.

Redl, C., Gfoehler, M. and Pandy, M.G. (2007) ‘Sensitivity of muscle force estimates to variations in muscle–tendon properties’, Human Movement Science, 26(2), pp. 306–319. Available at: 10.1016/j.humov.2007.01.008.

Riley, P.O., Krebs, D.E. and Popat, R.A. (1997) ‘Biomechanical analysis of failed sit-to-stand’, IEEE Transactions on Rehabilitation Engineering, 5(4), pp. 353–359. Available at: 10.1109/86.650289.

Riley, P.O., Schenkman, M.L., Mann, R.W. and Hodge, W.A. (1991) ‘Mechanics of a constrained chair-rise’, Journal of Biomechanics, 24(1), pp. 77–85. Available at: 10.1016/0021-9290(91)90328-K.

Roberts, T.J., Kram, R., Weyand, P.G. and Taylor, C.R. (1998) ‘Energetics of Bipedal Running: I. Metabolic Cost of Generating Force’, Journal of Experimental Biology, 201(19), pp. 2745–2751. Available at: 10.1242/jeb.201.19.2745.

Roebroeck, M.E., Doorenbosch, C.A.M., Harlaar, J., Jacobs, R. and Lankhorst, G.J. (1994) ‘Biomechanics and muscular activity during sit-to-stand transfer’, Clinical Biomechanics, 9(4), pp. 235–244. Available at: 10.1016/0268-0033(94)90004-3.

Rubenson, J., Lloyd, D.G., Besier, T.F., Heliams, D.B. and Fournier, P.A. (2007) ‘Running in ostriches (Struthio camelus): three-dimensional joint axes alignment and joint kinematics’, Journal of Experimental Biology, 210(14), pp. 2548–2562. Available at: 10.1242/jeb.02792.

Rubenson, J., Lloyd, D.G., Heliams, D.B., Besier, T.F. and Fournier, P.A. (2011) ‘Adaptations for economical bipedal running: the effect of limb structure on three-dimensional joint mechanics’, Journal of The Royal Society Interface, 8(58), pp. 740–755. Available at: 10.1098/rsif.2010.0466.

Savelberg, H.H.C.M., Fastenau, A., Willems, P.J.B. and Meijer, K. (2007) ‘The load/capacity ratio affects the sit-to-stand movement strategy’, *Clinical Biomechanics (Bristol*, Avon), 22(7), pp. 805–812. Available at: 10.1016/j.clinbiomech.2007.05.002.

Schaller, N.U., Herkner, B., Villa, R. and Aerts, P. (2009) ‘The intertarsal joint of the ostrich (Struthio camelus): Anatomical examination and function of passive structures in locomotion’, Journal of Anatomy, 214(6), pp. 830–847. Available at: 10.1111/j.1469-7580.2009.01083.x.

Schenkman, M., Berger, R.A., Riley, P.O., Mann, R.W. and Hodge, W.A. (1990) ‘Whole-Body Movements During Rising to Standing from Sitting’, Physical Therapy, 70(10), pp. 638–648. Available at: 10.1093/ptj/70.10.638.

Schultz, A.B., Alexander, N.B. and Ashton-Miller, J.A. (1992) ‘Biomechanical analyses of rising from a chair’, Journal of Biomechanics, 25(12), pp. 1383–1391. Available at: 10.1016/0021-9290(92)90052-3.

Scovil, C.Y. and Ronsky, J.L. (2006) ‘Sensitivity of a Hill-based muscle model to perturbations in model parameters’, Journal of Biomechanics, 39(11), pp. 2055–2063. Available at: 10.1016/j.jbiomech.2005.06.005.

Shia, V., Moore, T.Y., Holmes, P., Bajcsy, R. and Vasudevan, R. (2018) ‘Stability basin estimates fall risk from observed kinematics, demonstrated on the Sit-to-Stand task’, Journal of Biomechanics, 72, pp. 37–45. Available at: 10.1016/j.jbiomech.2018.02.022.

Sloot, L.H., Millard, M., Werner, C. and Mombaur, K. (2020) ‘Slow but Steady: Similar Sit-to-Stand Balance at Seat-Off in Older vs. Younger Adults’, Frontiers in Sports and Active Living, 2, p. 548174. Available at: 10.3389/fspor.2020.548174.

Smith, N.C. and Wilson, A.M. (2013) ‘Mechanical and energetic scaling relationships of running gait through ontogeny in the ostrich (Struthio camelus)’, Journal of Experimental Biology, 216(5), pp. 841–849. Available at: 10.1242/jeb.064691.

Smith, S.H.L., Reilly, P. and Bull, A.M.J. (2020) ‘A musculoskeletal modelling approach to explain sit-to-stand difficulties in older people due to changes in muscle recruitment and movement strategies’, Journal of Biomechanics, 98, p. 109451. Available at: 10.1016/j.jbiomech.2019.109451.

Thompson, L.A., Badache, M., Cale, S., Behera, L. and Zhang, N. (2017) ‘Balance Performance as Observed by Center-of-Pressure Parameter Characteristics in Male Soccer Athletes and Non-Athletes’, Sports, 5(4), p. 86. Available at: 10.3390/sports5040086.

Triviño, A., Davidson, C., Clements, D.N. and Ryan, J.M. (2023) ‘Objective comparison of a sit to stand test to the walk test for the identification of unilateral lameness caused by cranial cruciate ligament disease in dogs’, *Journal of Small Animal Practice*, p. jsap.13679. Available at: 10.1111/jsap.13679.

Yoshioka, S., Nagano, A., Hay, D.C. and Fukashiro, S. (2009) ‘Biomechanical analysis of the relation between movement time and joint moment development during a sit-to-stand task’, BioMedical Engineering OnLine, 8(1), p. 27. Available at: 10.1186/1475-925X-8-27.

Zajac, F.E. (1989) ‘Muscle and tendon: properties, models, scaling, and application to biomechanics and motor control’, Critical Reviews in Biomedical Engineering, 17(4), pp. 359–411.

